# Brain anatomy and molecular signaling predict neurophysiological dynamics across the lifespan

**DOI:** 10.64898/2026.04.03.715930

**Authors:** Christina Stier, Udo Dannlowski, Joachim Gross

## Abstract

Neural activity emerges from interactions between local cellular architecture, neuromodulatory systems, and large-scale cortical networks. Yet it remains unclear how this multiscale biological context constrains electrophysiological dynamics in humans and how this changes across the lifespan. We combined resting-state magnetoencephalography (MEG) from 350 adults (18–88 years) with cortical maps reflecting cytoarchitecture, myelination, metabolism, gene expression, and neurotransmitter receptors in a multivariate prediction framework. Specific markers explained most regional variance in MEG power spectra and temporal autocorrelation, similarly for both measures, revealing frequency- and timescale-specific signatures that followed canonical spectral boundaries. Age-related MEG patterns spatially aligned with markers of neuroinflammation, monoaminergic-cholinergic signaling, cortical development and myelination, and cerebrovascular organization. This work identifies key components of an anatomical and molecular scaffold and their relative importance for neural activity across the lifespan, informing future experimental perturbations and generative models.

## Introduction

The brain is organized across multiple spatial and temporal scales, forming the foundation for human cognition and behavior. Bringing together and understanding the relationship across these scales has been a longstanding question in neuroscience. Previous studies have delved into this problem from different angles, and many have used structural and functional MRI to link structural wiring to functional configurations (1–6). It emerged that structure-function coupling seems regionally heterogeneous, associations tend to be stronger in unimodal cortex and weaker in transmodal areas (7–12), and vary dynamically across time (13). Compared to BOLD signals measured in fMRI, the dynamics of neuronal population activity, as reflected in local field potentials (LFP) and therefore MEG/EEG signals, are arguably even more closely linked to and shaped by microstructural properties. The geometrical arrangement of neurons, such as their spatial alignment, directly affects the LFP, as does dendritic morphology (14). Neuronal population activity also differs across cortical layers, and electrophysiological signatures vary between cortical areas as a function of distinct laminar organisation. In addition, neuronal population dynamics are strongly shaped by gene expression, which regulates neurotransmitter receptors, transporters, and intracellular signaling cascades. Previous studies have linked genetic markers to specific brain rhythms (15,16), highlighting their heritable basis. Others have demonstrated that dynamic causal modelling of intracranial EEG data improves when normative receptor data are incorporated into the model (17).

At the macroscopic level, each cortical region comprises distinct neuron types, laminar organization, and connectivity that jointly influence the biophysics of electrophysiological signals. This endows each brain area with a characteristic functional fingerprint, allowing identification of brain areas based on the unique dynamic repertoire of their brain activity as represented in MEG signals (18). Interestingly, many microstructural features that shape brain activity change systematically across the cortex (19,20), establishing axes of organization such as gradients from unimodal to transmodal brain areas. These structural gradients are mirrored by corresponding functional gradients (21), which have been observed, for example, for neural power (22) and the frequency of the local dominant brain rhythms (23). In turn, the brain’s myeloarchitecture is spatially correlated with functional connectivity, particularly in the beta frequency range (24,25). Together, these studies demonstrate the intricate relationship between neuronal population dynamics and the underlying local microarchitectural scaffold. However, they have focused on a small set of functional, structural, or geometric measures, thereby limiting the development of integrated, multiscale model of biological interactions in the brain.

Recent efforts in collecting and merging PET- and MRI-derived data and forming large-scale consortia in recent years have opened up new possibilities for leveraging cellular and molecular metadata to refine models of structure-function relationships. This has resulted in a collection of brain maps that capture various aspects of microarchitecture, brain metabolism, cortical expansion, neurotransmitter receptors and transporters, and genomics (26,27). These resources have enabled several MEG studies linking local electrophysiological features to underlying biology. Shafiei and colleagues (28) used a multivariate prediction approach to link these maps to MEG activity based on data from the Human Connectome Project (HCP). Specifically, they characterized MEG signals in cortical parcels by 6,800 time-series signal features. Their analysis revealed that specific linear combinations of neurochemical and anatomical features predict corresponding combinations of electrophysiological features. However, the nature of specific feature-function associations remains unclear. Hansen and colleagues (29) used multiple linear regression to relate cortical MEG power in distinct frequency bands to specific neurotransmitter receptor densities. Based on average MEG data from 33 young adults, they found close correspondence between mostly inhibitory receptor distributions and power in canonical frequency bands, except for high-gamma (29). Extending this work, Siebenbühner et al. (30) have provided evidence that local receptor architecture shapes MEG networks in a spectrally specific manner in average data from 70 participants. The results differed for the MEG connectivity measure and frequencies investigated. While these studies provide important groundwork for understanding structure–function relationships, their correlational nature limits insight into predictive capacity, and the specificity of feature–function associations remains incompletely understood.

Lifespan change adds another critical dimension to the investigation of structure-function coupling. The brain undergoes continuous reorganization throughout adulthood, characterized by systematic changes in structural organization, molecular composition, and functional activity. fMRI studies spanning young adulthood have demonstrated coordinated reorganization of functional networks alongside structural wiring changes (31) and have identified how structure-function coupling in prefrontal regions supports age-related improvements in executive function (8). Work covering an age range up until old age found overall reduced structure-function coupling with age, particularly in sensorimotor regions (32). Others have described an interplay between declining functional segregation and age-related changes in white matter integrity (33). However, similar efforts linking electrophysiological aging signatures to concurrent structural and molecular changes covering brain dynamics at much higher temporal resolution have been scarce (34–37). Age trajectories for delta and beta oscillations have spatially co-localized with those for cortical thickness (38), indicating structure-function relationships across the lifespan.

Moreover, when studying age effects on brain activity at the millisecond scale, most literature has focused on signal amplitude, such as power, with resting-state eyes-closed spectral power decreasing in low frequencies and increasing in higher frequencies across healthy aging (39–44), with prominent effects observed in frontal and temporal cortex (38,45–47). In contrast, age-related changes in the temporal structure of brain activity have been less thoroughly explored (21). However, measures such as autocorrelation (AC) have been effective indices of age-related changes in the brain connectome (48) and cortical hierarchy (22). Using MEG, we recently demonstrated that temporal autocorrelation (AC) patterns, particularly within visual and temporal cortices, provided a robust marker for chronological age and outperformed traditional spectral measures in predicting age (49). Collectively, these findings motivate investigations of how structure-function relationships manifest across the lifespan in both spectral and temporal measures, probing biological substrates of neural aging at the millisecond timescale.

In summary, it is well established that electrophysiological brain activity is significantly shaped by underlying microarchitecture and that structure, function, and presumably also their coupling change across the lifespan. However, the unique contributions of individual structural and neurochemical features to regional brain dynamics, and their relative importance for explaining inter-regional variability, remain unclear. Specifically, it has been unknown what part of the variance in regional brain activity can be explained by such features, and whether this explanatory power changes across the lifespan. Here, we address these fundamental questions by constructing a cross-validated, multivariate predictive model that links multiscale anatomical and molecular features to well-studied parcel-wise descriptors of resting-state MEG in a large adult lifespan cohort (n = 350, (50,51)). Importantly, several brain rhythms occur simultaneously, orchestrating brain activity in a coordinated manner. Hence, we aimed at predicting the full electrophysiological spectrum and temporal structure in a multivariate manner, rather than isolated band-limited measures, using partial least squares regression. With this framework, combined with incremental feature selection, we can quantify how much variance in regional MEG dynamics (individual fingerprints of brain areas) can be explained by local structural and neurochemical features. The model allows us to characterize the individual importance of the features and the aspects of spectral and temporal dynamics they modulate. Finally, we capitalize on the large adult lifespan cohort by investigating changes in this predictive model across age. We found that structural and neurochemical features jointly explain a substantial proportion of regional variance in MEG power spectra and temporal autocorrelation across the cortex (average R^2^ of ∼0.8). Second, explanatory power varied markedly across features but was similar for power and autocorrelation. Third, single features showed frequency-specific positive and negative associations with spectral power that followed canonical frequency boundaries (for example, positive effects on alpha band power and negative effects on gamma power). Fourth, the spatial distribution of specific features significantly explained age-related changes in MEG power and autocorrelation. Together, this work delineates aspects of brain microstructure and neurochemistry that constitute the backbone for cortical electrophysiological dynamics across the adult lifespan.

## Results

### Prediction of regional differences in spectral power

We initially predicted the full parcel-wise, group-averaged power spectrum from a comprehensive set of brain maps depicting neurotransmitter receptor and transporter distributions, metabolism, gene expression, and microstructure. In a cross-validated scheme, we repeatedly fit partial least squares regression (PLSR) in 180 brain parcels and tested in 20 held-out parcels. Before the fold split, we normalized the power spectrum by subtracting the global spectral mean of each parcel. This centers the spectrum around zero and removes variance that is due to the background activity (1/f structure) observed in the power spectrum. Within each fold and frequency bin, we then z-scored across parcels (see Methods). Thus, the model learns to predict relative deviations from the mean across parcels rather than absolute power. In other words, we aimed at explaining the differences between parcels. We evaluated the model’s performance by quantifying the explained variance relative to simply predicting the mean (predicted R^2^) and testing its significance against a null distribution generated using random spherical rotations and re-assignment of the data (spin-test (93)). We further estimated the median identifiability of individual brain parcels (see Methods), which would lead to a rank of 200 given a perfect model. Our model predicted regional power differences with high accuracy across all parcels (R^2^ = 0.83, p_spin_ < 0.05, median rank 195). To identify which predictors contributed most to the fit, we computed variable importance in projection (VIP) scores. **Figure 2** shows that VIP scores varied greatly across maps. Overall neuronal density (cyto-intensity) was by far the best predictor. This is followed by the mu-opioid receptor density map (MOR) and the first principal component of gene expression (genepc1). Amongst the 24 predictors with a VIP score above 1, six are measures describing the brain’s cyto- and myeloarchitecture, eleven represent neurotransmitter receptor distributions, and other single measures depict gene expression, spatial organization including gradients and cortical hubness, metabolism, and neurotransmitter transporters and enzymes.

**Fig. 1.**
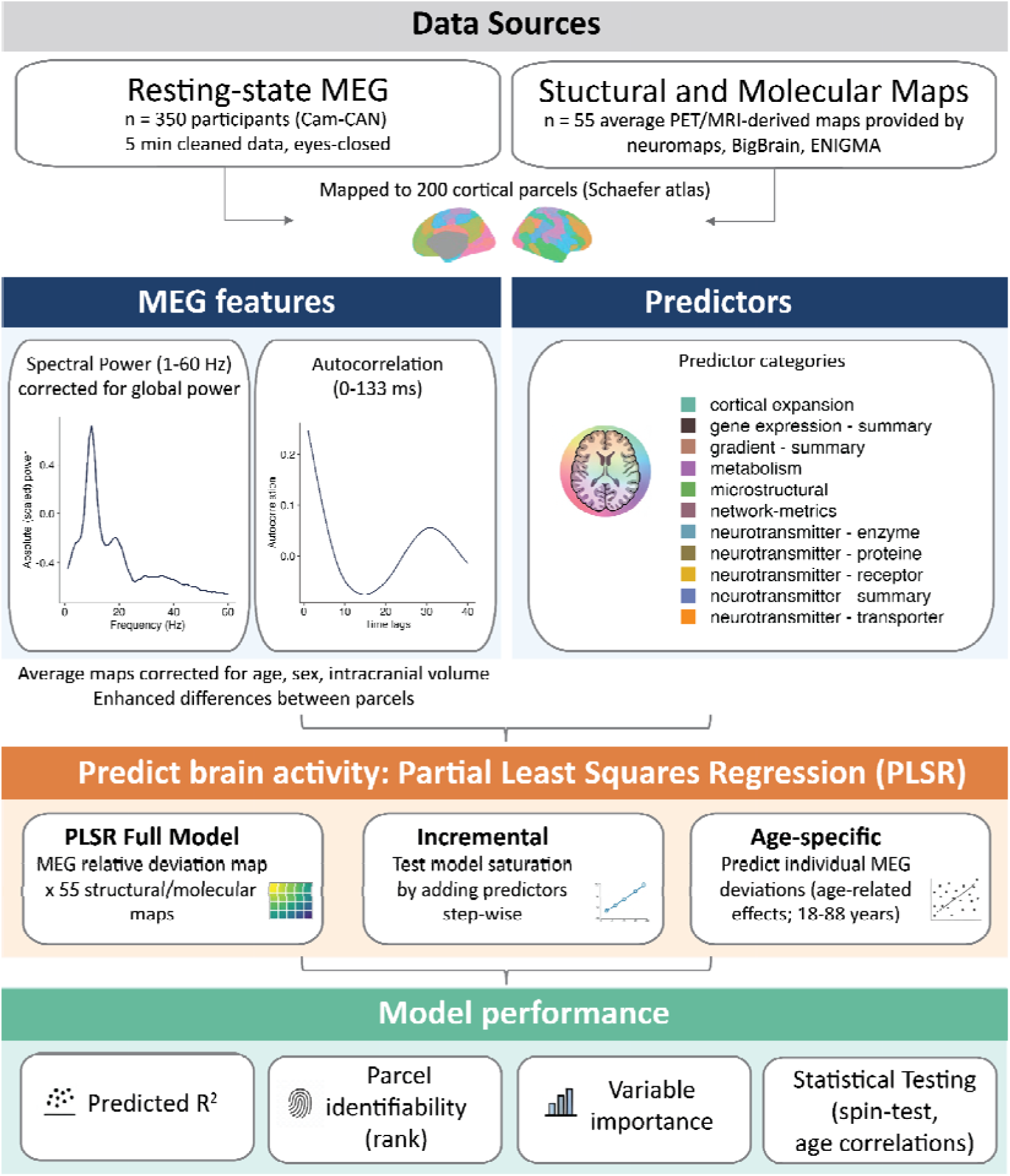
Overview of data sources, feature extraction, and modelling pipeline. We used resting-state MEG recordings from 350 healthy adults (18–88 years) provided by the Cambridge Center for Aging and Neuroscience (Cam-CAN) and 55 population-average structural and neurochemical brain maps derived from PET/MRI resources (neuromaps (26), BigBrain (52), ENIGMA (27)), all parcellated into 200 cortical regions using the Schaefer atlas (53). From the source-projected parcel-wise MEG time series, we computed power spectra (1–60 Hz) corrected for global broadband power and temporal autocorrelation functions (0–133 ms). We generated average maps of these features after regressing out age, sex, and intracranial volume. Structural and neurochemical predictors are listed in detail in Table 1 and summarized here in respective categories. We used partial least squares regression (PLSR) to predict regional (parcel-wise) differences in MEG power spectra and autocorrelation from the predictor set, implementing cross-validated full models, and an incremental procedure to assess how many and which set of predictors achieve similarly high prediction accuracy. We then built age-specific models, relating subject-level MEG deviations from the group average, reflecting age effects in the MEG metrics, to the same predictor maps. Model performance was quantified using cross-validated prediction R^2^, parcel identifiability (median rank of the correlation between observed and predicted parcel-wise profiles), variable importance in projection (VIP) scores to rank predictors, and spatial spin tests and age correlations to assess statistical significance.

**Fig. 2.**
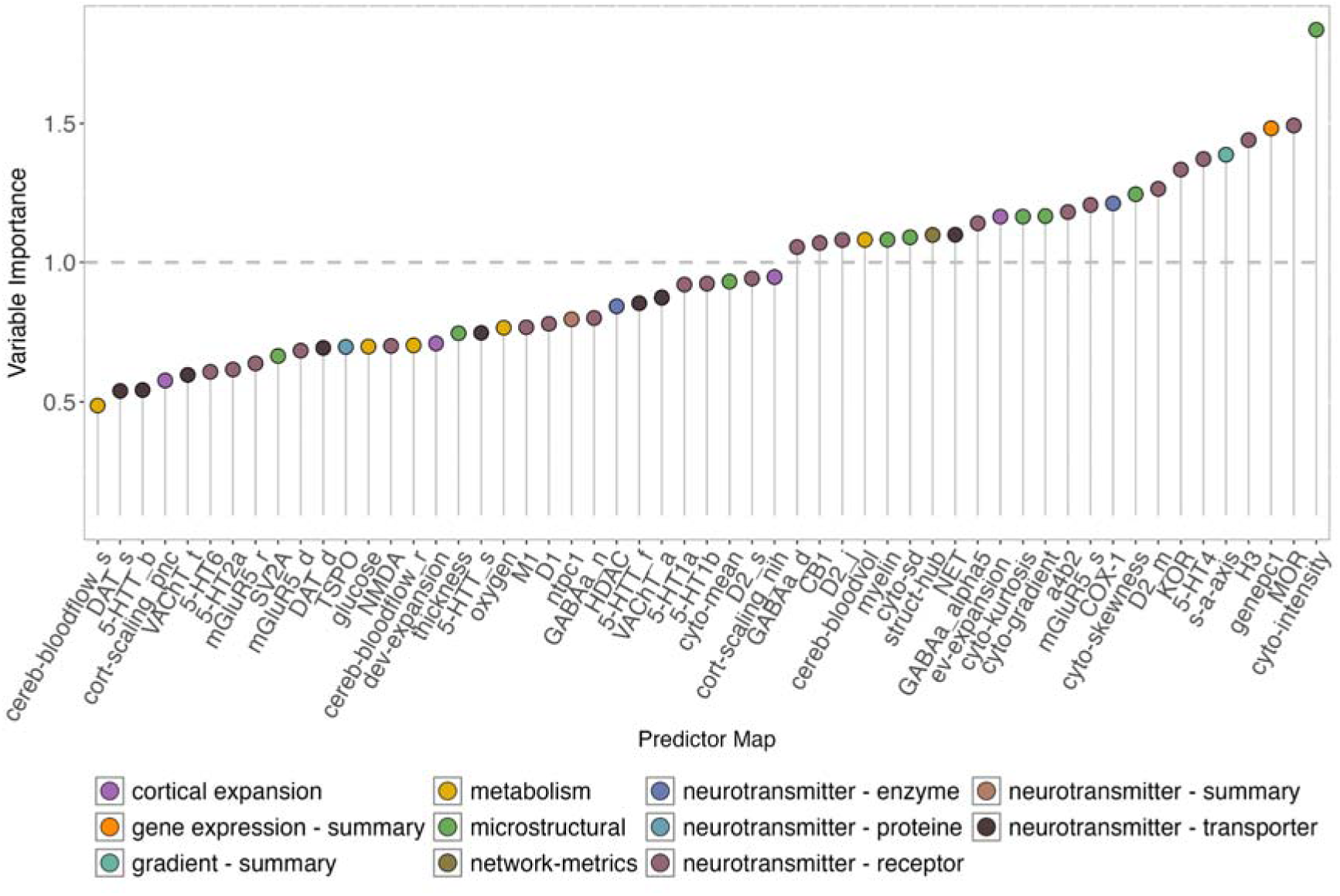
Importance of single features predicting regional power differences. Each dot indicates the importance of a single structural or neurochemical feature, with colors indicating the respective category. Features with a variable importance score larger than 1 (orange dashed line) are considered key contributors to the partial least squares regression model predicting parcel-wise power differences based on 55 predictor maps (features). See Table 1 for feature abbreviations.

**Table 1.**
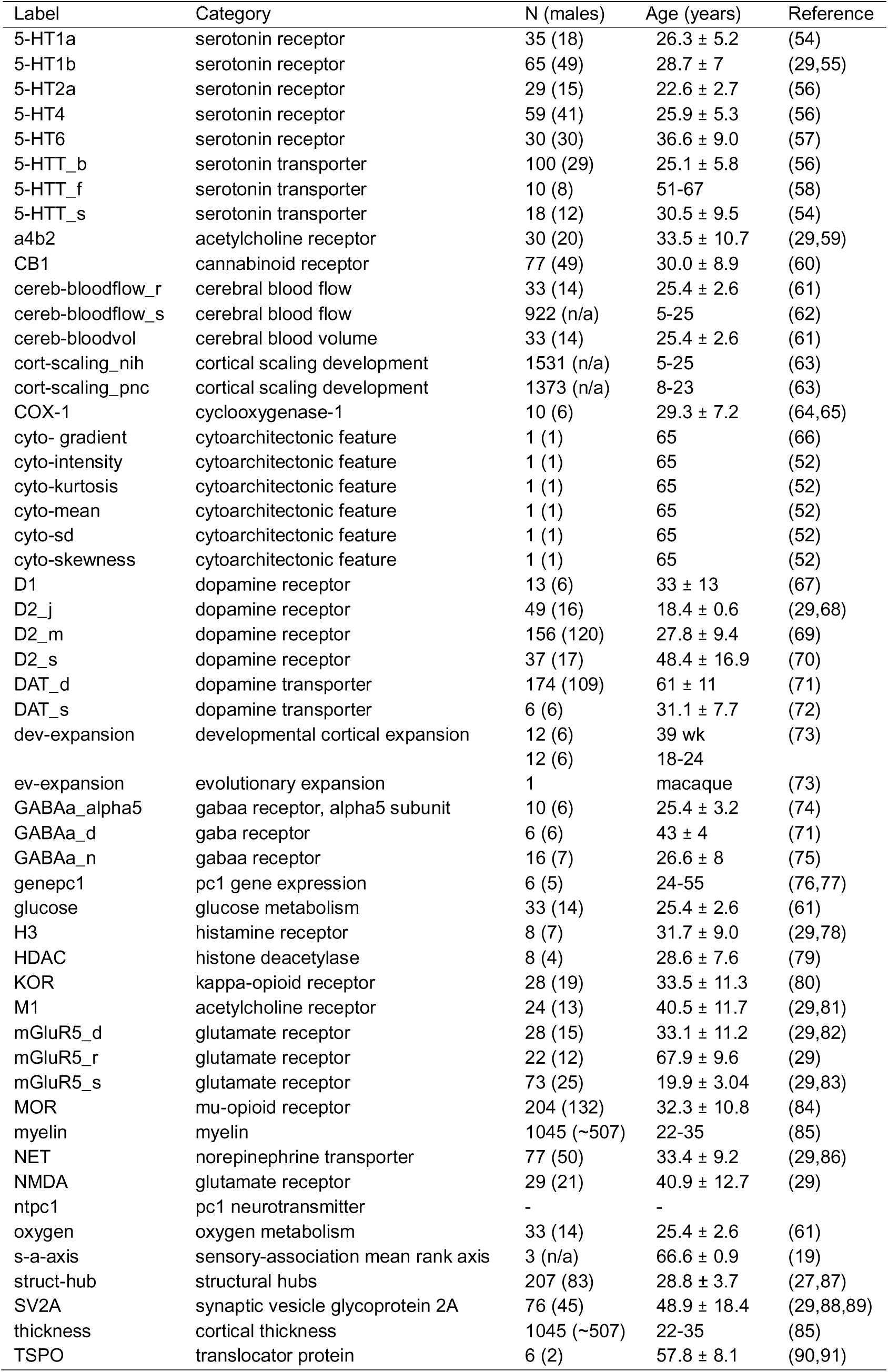
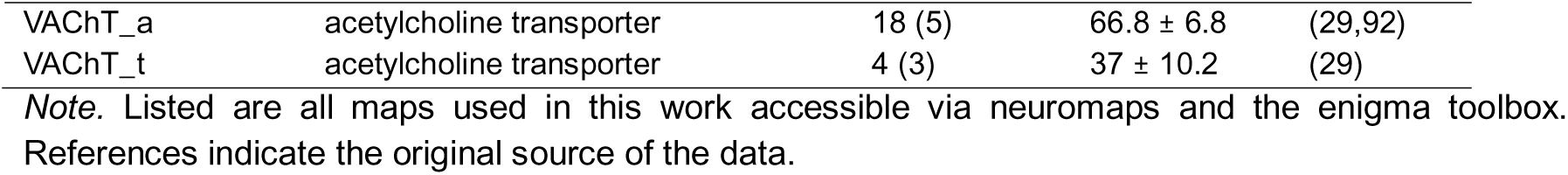
Overview of structural and neurochemical predictors.

Next, we studied the specific effects that individual maps have on spectral power. **Figure 3** shows the regression weights from the PLSR model for each map and frequency. Interestingly, individual maps exhibit frequency-specific, reversed relationships to spectral power. For example, overall neuronal density (cyto-intensity), the first principal component of gene expression (genepc1), and the serotonin receptor 5-HT4 were positively associated with MEG power differences in the alpha frequency range, but negatively in low (< 7 Hz) or high (> 12 Hz) frequencies. Other highly contributing maps, such as opioid (MOR, KOR) and histamine (H3) receptors and cortical hierarchy (s-a-axis), were associated in opposite directions: positively with low (< 7 Hz) and high frequencies (> 12 Hz), and negatively with alpha power. Prediction weights for other brain maps were smaller and less distinct in their prediction patterns. However, a few further maps shall be mentioned due to their specific profiles: evolutionary expansion of the human brain (ev-expansion), a cytoarchitectural gradient (sensory-fugal), and cerebral blood volume (cereb-bloodvol) showed large model weights in the beta frequency ranges, pointing to strong (anti-)correlations with MEG power differences.

**Fig. 3.**
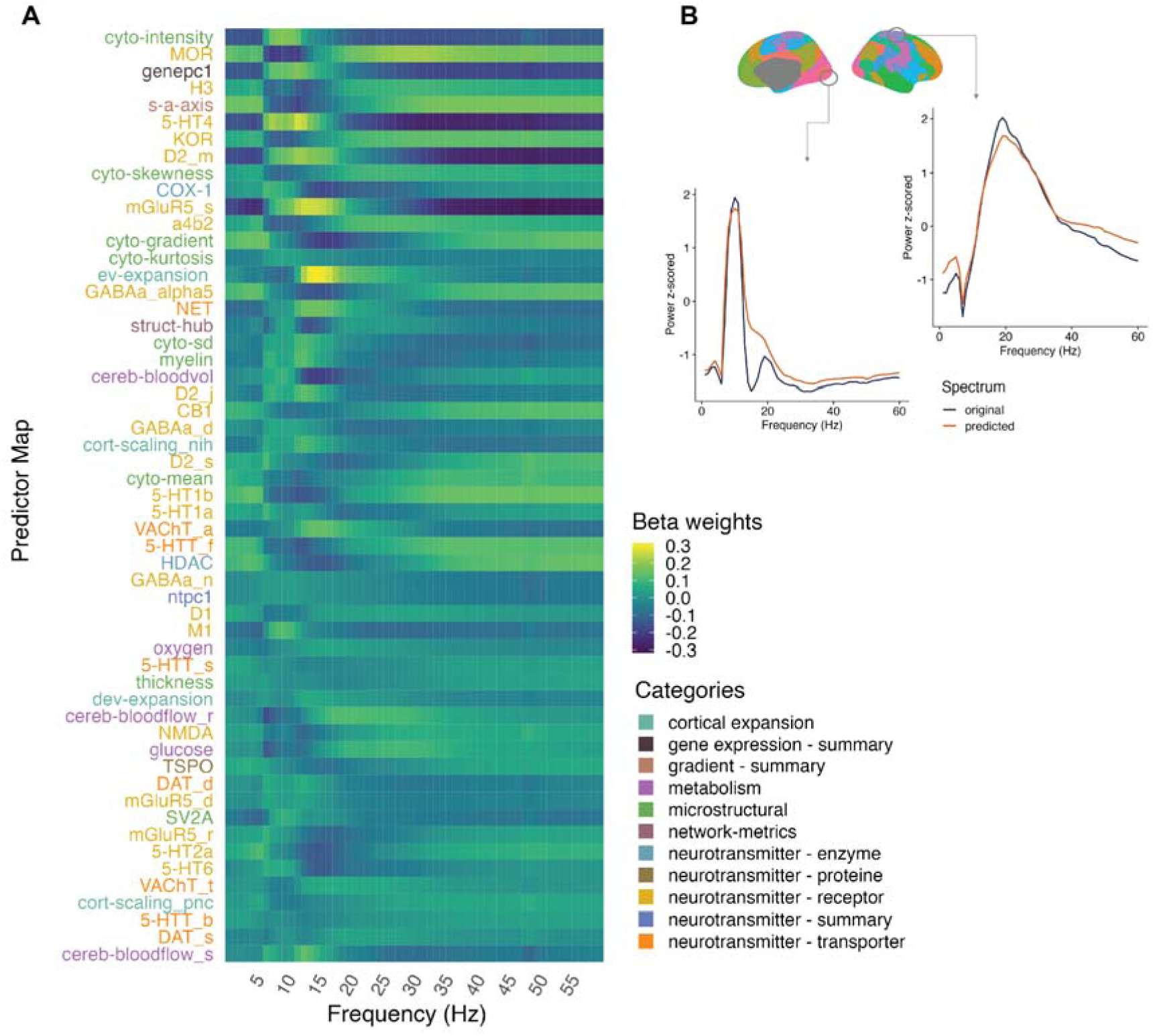
Prediction outcome for regional power differences. A) This heatmap shows partial least squares regression weights for each of the predictor maps (n=55). The respective maps are ranked according to their average variable importance and indicate positive versus negative relationships between the predictors and the regional power spectra (1-60 Hz). For visualization purposes, the labels are colored according to their predictor category. Key predictors include overall neuronal density (cyto-intensity), opioid (MOR), histamine (H3), dopamine and serotonin receptors, and the first principal component of gene expression (genepc1). B) Example plots showing the original and predicted spectrum within single parcels of the visual (left) and somatomotor (right) cortex based on all predictor maps. Note that the prediction accuracy was computed across all parcels, yielding a predicted R^2^ of 0.83.

Next, we quantified the number of maps required to achieve prediction performance equivalent to that of the full set of maps by incrementally adding predictors. An R^2^ of 0.8 was achieved using 18 brain maps, based on their single or additive contribution to the prediction of average MEG power (**Figure 4**). The key predictors in the incremental analysis were similar to those from the full-model VIP ranking, although their order shifted slightly, likely because the number of PLS components was increased iteratively during the selection procedure. Overall, a modest subset of maps provides most of the predictive power for parcel-wise spectral profiles.

**Fig. 4.**
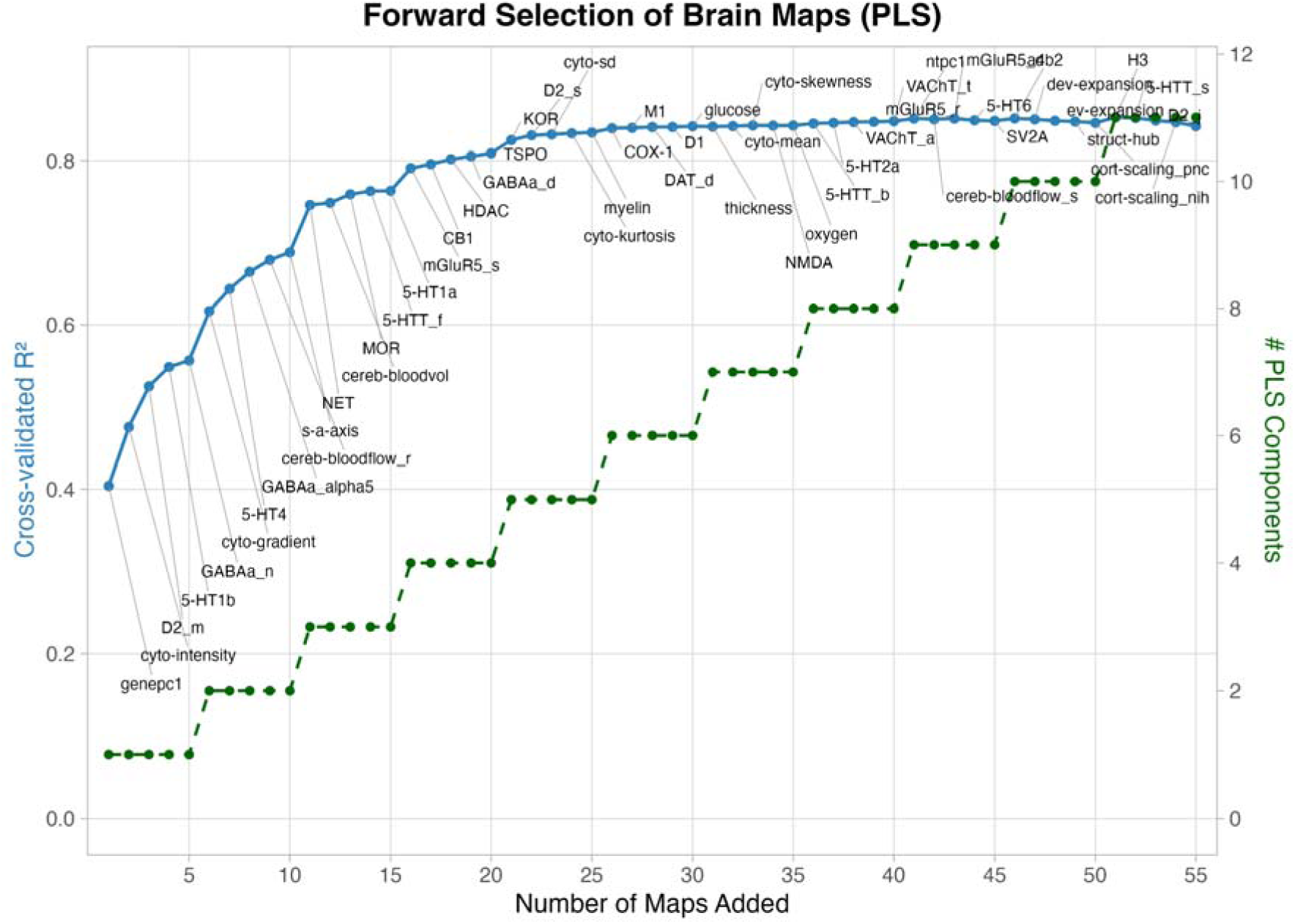
Incremental prediction of regional power differences. The blue trajectory in the plot represents the absolute cross-validated R^2^ after adding the respective brain map to the partial least squares regression model. We first evaluated which single map performed best and included another map, bringing the R^2^ up to the next highest. In parallel, we gradually increased the number of latent components, that is, after every fifth predictor (green trajectory).

### Prediction of regional differences in temporal autocorrelation

As outlined earlier, measures capturing the temporal structure of MEG signals have been a robust indicator of the brain’s hierarchical organization (22) and chronological age (22,49). We therefore repeated the previous analyses using the temporal autocorrelation (AC) function instead of the power spectrum. AC was computed between signals across 40 time lags, corresponding to shifts of 3.3 ms up to 133 ms in our data. As for power, we predicted relative deviations from the mean across parcels. This yielded a similarly high prediction accuracy at the level of brain parcels compared to power (R^2^ = 0.88, p_spin_ < 0.05, median rank 193). There was a considerable overlap between predictors of power and AC: seven of ten top-ranking maps were shared (**Figure 5**). Compared to the results for power, one further map shows a slightly higher ranking or prominence for autocorrelation in both the full model and the reduced feature model, namely acetylcholine receptor distribution (‘a4b2’). Notably, fewer maps were needed to reach equivalent performance for AC than for power: only 12 predictors sufficed to achieve a similar R² (**Figure 6**).

**Fig. 5.**
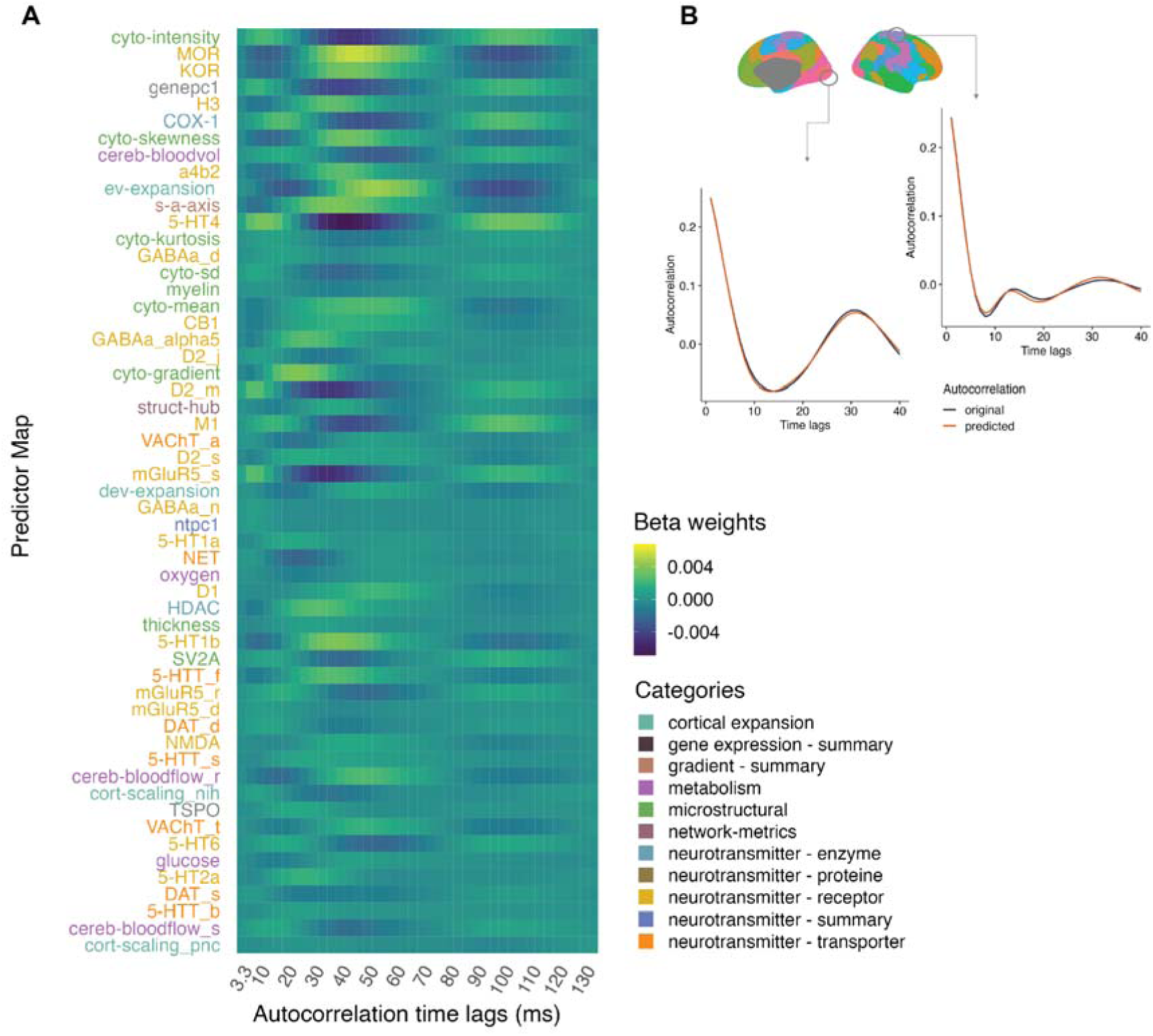
Prediction outcome for temporal autocorrelation. A) This heatmap shows partial least squares regression weights for each of the predictor maps (n=55). The respective maps are ranked according to their average variable importance and indicate positive versus negative relationships between the predictors and the regional power spectra (1-60 Hz). For visualization purposes, the labels are colored according to their predictor category. B) Example plots showing the original and predicted autocorrelation function within single parcels of the visual (left) and somatomotor (right) cortex based on all predictor maps. Note that the prediction accuracy was computed across all parcels, yielding a predicted R^2^ of 0.88.

**Fig. 6.**
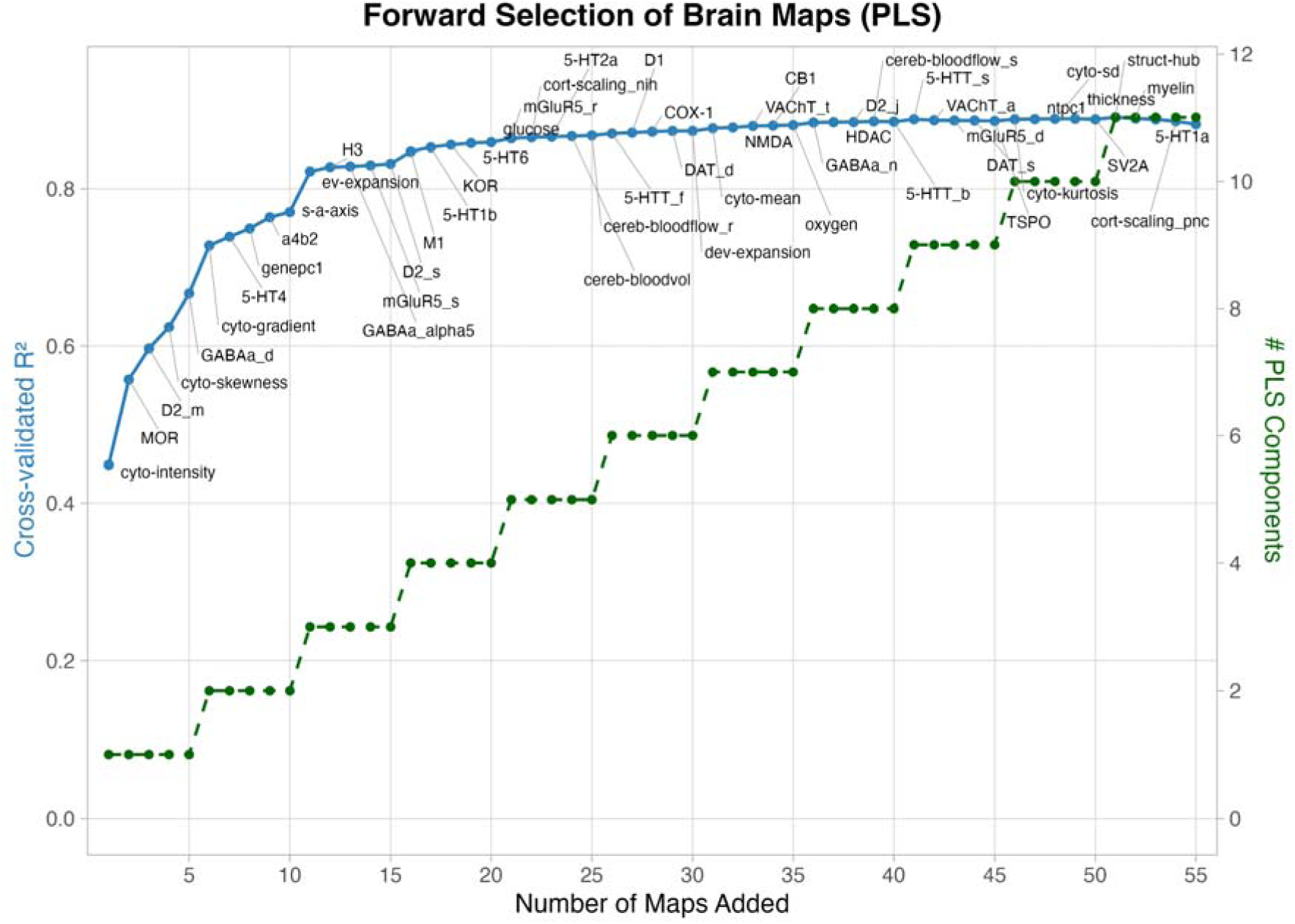
Incremental prediction of regional differences in temporal autocorrelation. The blue trajectory in the plot represents the absolute cross-validated R^2^ after adding the respective brain map to the partial least squares regression model. We first evaluated which single map performed best, and included another map, adding up to the next highest R^2^. In parallel, we gradually increased the number of latent components, that is, after every fifth predictor (green trajectory).

### Mapping of neurophysiological aging

Moreover, we investigated which predictors reflect age-related variation in MEG metrics. Leveraging subject-level data from the Cam-CAN set (n=350, 18-88 years), we computed each individual’s parcel-wise deviations from the covariate-adjusted average map. For each subject and metric (power or AC), we fitted a PLSR model predicting these parcel-wise deviations using the 55 structural and neurochemical maps. This yielded subject-specific model performance and predictor weights. For power, individual predictions (R²) ranged from 0.66 to 0.97 (**Figure 7**). To get an estimate of which features reflect neurophysiological aging, we correlated chronological age with the weights for each predictor. **Figure 8** shows the correlation coefficients (r) for each predictor map and frequency bin. Again, the maps and age-related neurophysiological deviations co-localized differently across the frequency spectrum, yielding positive and negative associations. The strongest average associations (mean |r| > 0.3 across frequencies) were found for the cyclooxygenase (COX-1), myelin, serotonin (5-HT1a), evolutionary expansion (ev-expansion), and histamine (H3) maps.

**Fig. 7.**
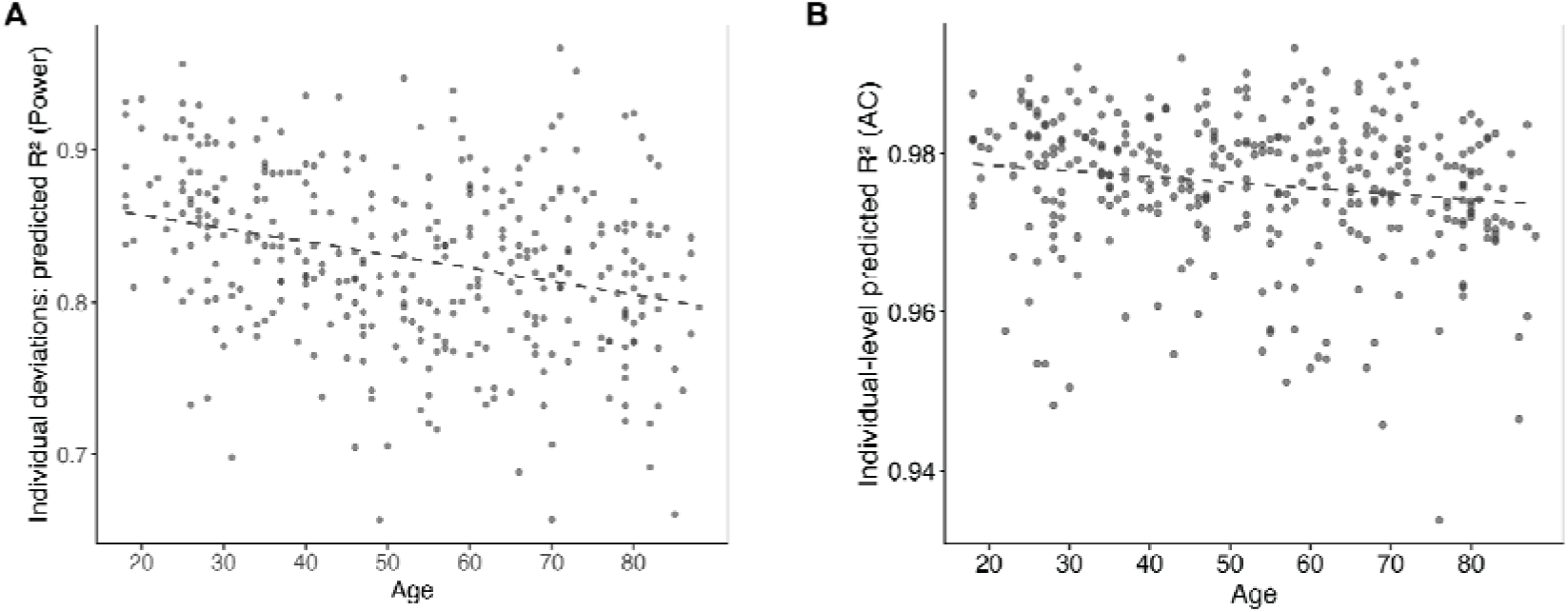
Prediction of power and autocorrelation patterns across the lifespan. Each dot represents the predicted R^2^ for each individual from the sample (n=350, 18-88 years) and quantifies to what extent individual deviations from the sample’s average were predicted based on the full set of structural and neurochemical predictors (n=55). Given the broad and balanced age distribution of the sample, individual deviations from the average reflect age-specific effects in the MEG data.

**Fig. 8.**
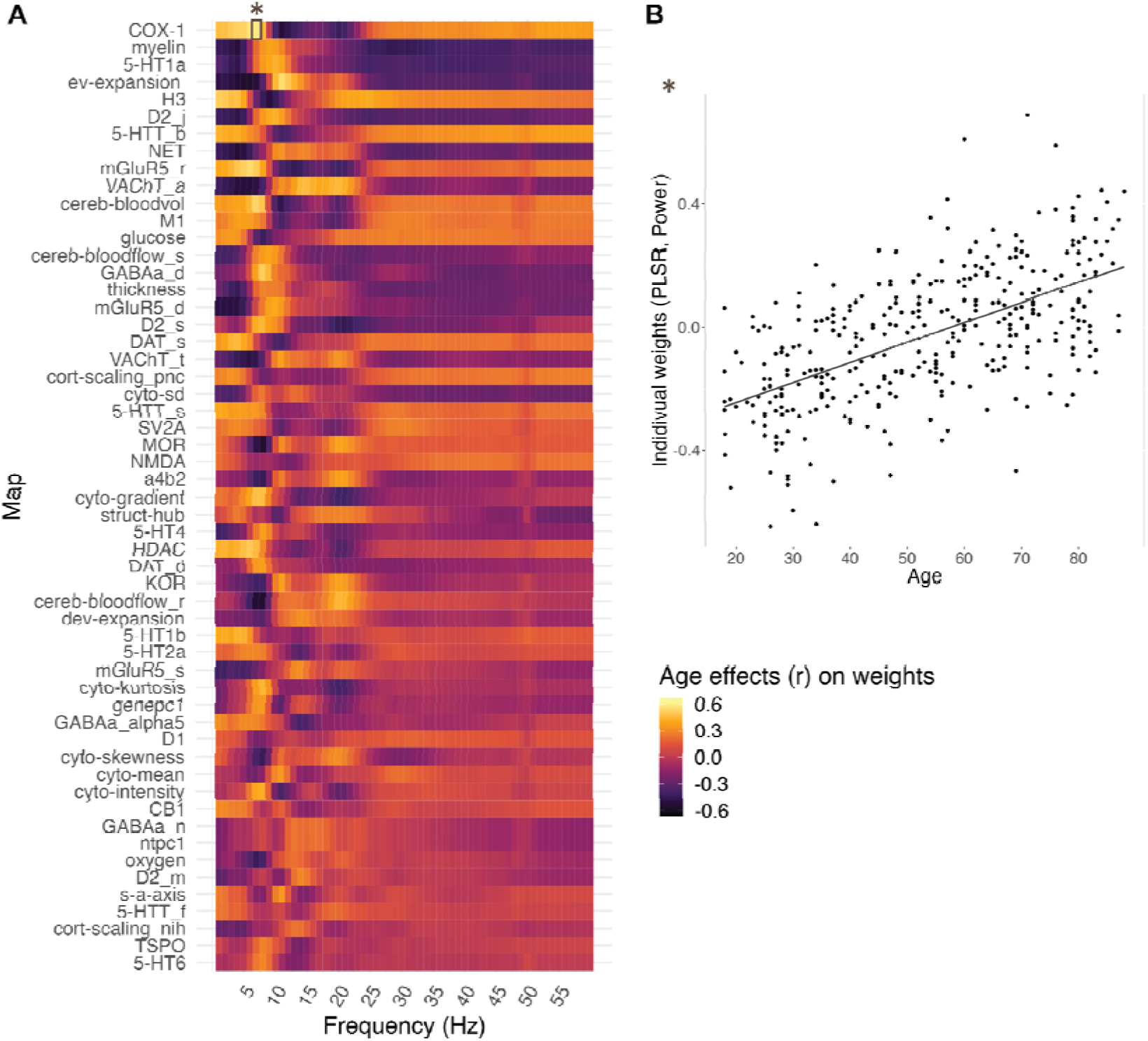
Structural and neurochemical maps reflecting aging effects in MEG power. A) This heatmap shows Pearson correlation coefficients between individual PLS model weights and chronological age (n=350, ages 18-88 years) for all 55 predictor maps. Maps are ranked by average correlation across the frequency spectrum. Color-coding indicates positive (warm) and negative (cool) associations with aging. B) An example plot for the relationship between subject-level prediction weights and age, in this case for cyclooxygenase (COX-1) and MEG power differences at 7 Hz. COX-1 showed the strongest mean age effects across the frequency spectrum, with a positive association between individual weights and age. PLSR = partial least squares regression

For AC, individual predictions (R²) were substantially more accurate than for power, ranging from 0.93 to 0.99 (**Figure 9**). Approximately half of the top ten maps for AC overlapped with power, such as cyclooxygenase (COX-1) and evolutionary expansion (ev-expansion), with the most prominent associations with age (mean |r| > 0.3 across time-lags), besides serotonin (5-HT1a) and acetylcholine transporter (VAChT) maps. Additionally, maps of cerebral bloodflow and opioid receptors (MOR) were identified as particularly relevant for age effects in temporal autocorrelation (mean |r| > 0.3).

**Fig. 9.**
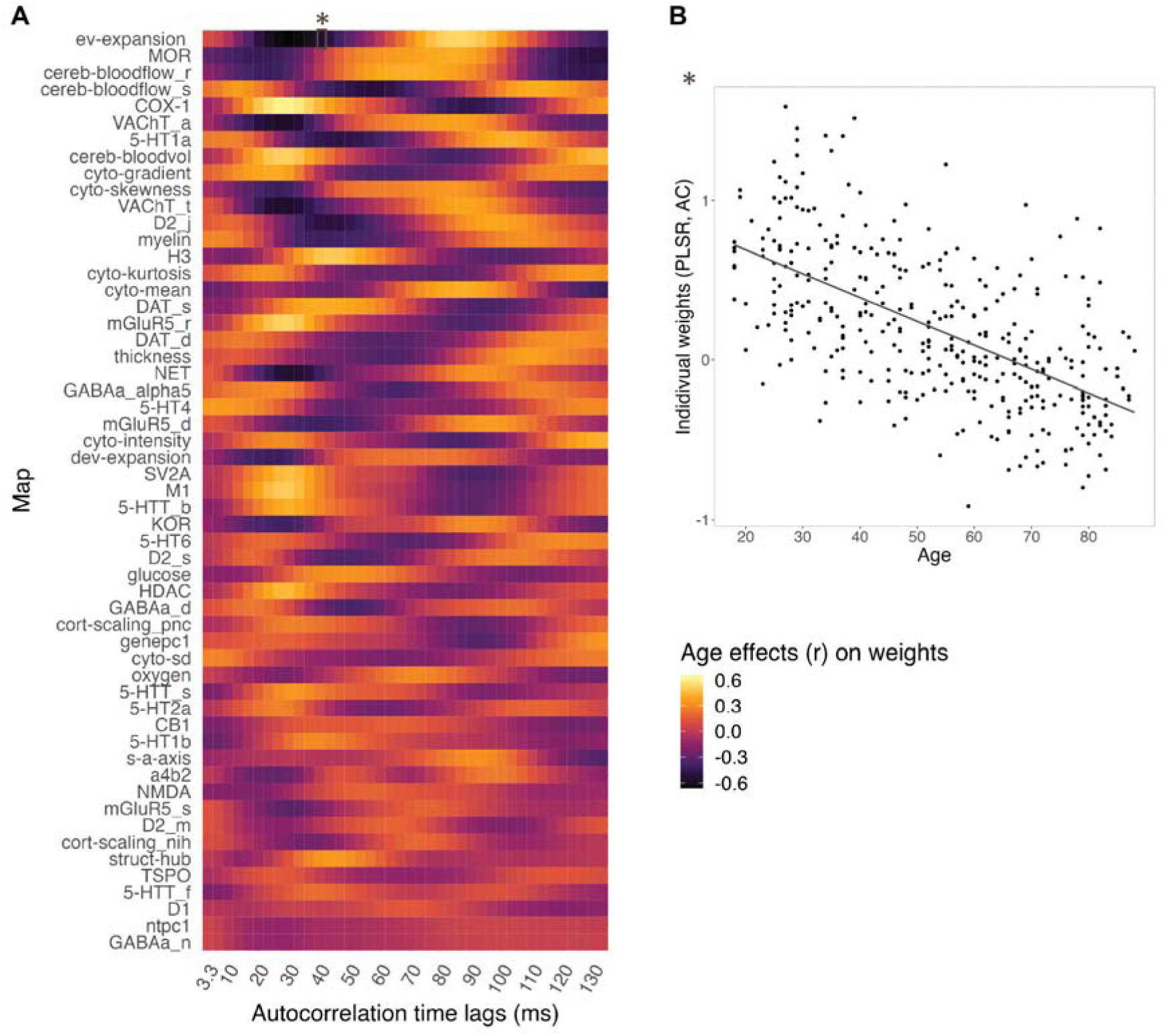
Structural and neurochemical maps reflecting aging effects in the MEG autocorrelation function. A) Heatmap showing Pearson correlation coefficients between individual PLS model weights and chronological age (n=350, ages 18-88 years) for all 55 predictor maps. Maps are ranked by average correlation across time lags. Color-coding indicates positive (warm) and negative (cool) associations with aging. B) An example for the relationship between subject-level prediction weights and age, in this case for the evolutionary expansion map (ev-expansion) and differences in temporal autocorrelation at ∼36 ms. Ev-expansion showed the strongest age effects across the time lags, with a negative association between individual weights and age. PLSR = partial least squares regression, AC = Autocorrelation

## Discussion

In this study, we show that regional resting-state electrophysiological signatures, both power spectra and temporal autocorrelation functions, can be robustly predicted from a comprehensive set of structural and molecular cortical maps. Using multivariate prediction at the parcel level, we find that a small subset of features including neuronal density, a dominant axis of cortical gene expression, and several neurotransmitter receptor maps explain a substantial portion of the spatial variance in electrophysiological measures. Temporal autocorrelation was even more strongly constrained by these features than spectral power and required fewer predictors to achieve comparable performance. Finally, age-related deviations in neurophysiological activity were systematically aligned with markers related to neuroinflammation, monoaminergic and cholinergic neurotransmission, cortical myelination, and vascular organization.

Our findings extend prior work linking MEG measures to receptor and structural maps by focusing on the identifiability of parcelwise activity patterns using linear prediction rather than simple spatial correlation. This approach allowed us to quantify the relative importance of different feature classes, which varied substantially. Core structural components, most prominently neuronal cell density and overall volume of stained somata, ranked highest for both spectral power and temporal autocorrelation, in line with earlier reports using average MEG data and multivariate prediction approaches (28). This finding is conceptually intuitive, as neuronal populations constitute the primary generators of the electromagnetic fields measured by MEG. Regions with higher neuronal density may support stronger local field potentials, thereby shaping both the amplitude and temporal structure of neural signals. Interestingly, increases in neuronal spiking were associated with increases in gamma and theta power and decreases in alpha/beta power (94). This precisely matches the spectral pattern observed in the regression weights in **Figure 3** for cytoarchitectural properties (and other maps), showing how a generic dynamic brain motif is regulated by local microarchitecture. This motif is well-known and strongly expressed in the sensory cortex during sensory stimulation or in the motor cortex during movement and is reflected in the increase of low-frequency and gamma power and the suppression of alpha/beta rhythms (95–98). The strong posterior–anterior gradient of neuronal density further mirrors well-established macroscale organizational axes of the cortex. This aligns with earlier observations that fundamental architectural gradients provide a scaffold for regional differences in resting-state dynamics (22,23). Similarly, large-scale gene expression patterns were highly informative, summarized here by the first principal component across thousands of genes, presumably reflecting major cell classes (76,99,100) and more broadly, laminar architecture (76,99). In addition to features strongly aligning with a posterior-anterior organization in the brain, an archetypal map of intrinsic spatial organization along a sensorimotor-to-association-axis (s-a-axis) (19) was indicative of MEG activity, albeit less pronounced. Hence, our study’s results synthesize existing observations on structure-function coupling and give insights into which markers constrain neural dynamics along the full spectral profile and temporal dependencies.

Beyond microstructural features, several receptor maps with predominantly inhibitory modulatory effects, most prominently opioid (MOR, KOR), histamine (H3), and dopamine (D2), exhibited strong, frequency-specific associations with MEG power and autocorrelation. Other high-ranking candidates included mixed serotonin (5-HT) subtypes and acetylcholine-related markers (a4b2), typically associated with net excitatory effects on brain circuits. These findings align with previous reports of spatial correspondence between receptor distributions and oscillatory power in selected frequency bands, including strong associations of theta–alpha and low gamma power with MOR, H3, and α4β2 maps (29). We extend these investigations in important ways. First, by modeling the entire power spectrum simultaneously, we characterized the direction of association between each predictor map and specific frequencies. For example, the MOR receptor, amongst our most predictive features, is predominantly expressed in interneurons and involved in regulating inhibitory signal transmission (101,102). Consequently, its regression weights show the opposite correlation to the spectral motif earlier discussed for neuronal density (i.e., positive associations with alpha power and negative associations with slower and faster rhythms). Second, in contrast to earlier work based on relatively small or age restricted samples, our average MEG map was derived from 350 individuals spanning adulthood, with age, sex, and intracranial volume residualized, thereby providing a robust reference. Third, by incorporating temporal autocorrelation, we show that similar receptor systems also constrain intrinsic timescales of neural activity.

The close correspondence between receptor profiles and temporal autocorrelation is notable given the mathematical link between signal power and autocorrelation via the Wiener–Khinchin theorem (103). However, fewer maps were required to predict autocorrelation at a high level of accuracy, suggesting that time-series characteristics may provide a more parsimonious readout. This resonates with recent work showing that time-domain metrics can outperform spectral features (22,104), particularly when certain assumptions are violated. Brain activity is often non-sinusoidal or arrhythmic, posing methodological challenges for estimating spectral features using Fourier methods. Instead, the AC function reflects the specific waveform shape and is normalized between -1 and 1. Our results therefore support the view that temporal autocorrelation is a robust marker for linking molecular architecture to macroscopic brain activity, particularly in the context of healthy aging (49).

To probe age-specific effects, we did not explicitly model interactions between age and structure-function coupling parameters. Instead, we computed each individual’s deviation from the covariate-adjusted group-average power spectrum and autocorrelation function, and related these subject specific residuals to the full set of structural and molecular maps. This revealed that age related deviations in MEG dynamics were best captured by markers of neuroinflammation, neuromodulation, cortical myelination, and vascular organization. COXL1 topography, in particular, emerged as a robust predictor of aging effects for both power spectra and autocorrelation functions. COX-1 enzymes are responsible for homeostatic prostaglandin synthesis(105)(105) and have been implicated in neurodegeneration(106)(106) and Alzheimer’s disease through inflammation(107)(107). MEG age effects also co-localized with monoaminergic systems, including serotonin receptor binding profiles and dopaminergic markers, and, for MEG power, with histaminergic maps. Vesicular acetylcholine transporters similarly emerged as important. These systems are known to interact through shared circuits (108,109). For example, basal forebrain cholinergic projections modulate cortical and hippocampal pyramidal neurons, and corticostriatal glutamatergic inputs converge with nigrostriatal dopaminergic projections in the striatum to shape core cognitive and motor functions (109). Thus, cholinergic pathways play an important role in memory (110,111) and in neurodegenerative disorders such as Alzheimer’s disease (112) and Parkinson’s disease (113,114). Our findings therefore dovetail with evidence suggesting that the aforementioned neuromodulatory systems are involved in age-related disorders and cognitive (dys-)functions, as well as with normative modeling studies showing spatial overlap between functional brain aging and these receptor distributions (115).

Beyond neurochemical systems, both MEG markers showed age-related variation along evolutionary expansion patterns. These closely resemble maps of postnatal cortical surface-area expansion (73), reflecting protracted maturation in respective areas such as higher-order association cortex. Several additional structural markers exhibited modality-specific associations with aging. Intracortical myelin, indexed by the T1-to-T2-weighted ratio, was particularly informative for age effects in the power spectrum, with opposite effects in low and high frequency bands relative to alpha/low-beta range. This pattern is consistent with prior work showing that cortical myelin follows an inverted U shaped lifespan trajectory (116) and that myelin changes progress along a posterior–anterior gradient(117)(117), potentially contributing to previously reported age related shifts in functional brain activity (22,118). In contrast, vascular markers such as cerebral blood flow and blood volume were more strongly associated with age-related deviations in the temporal autocorrelation function, aligning with evidence that cerebral perfusion and neurovascular coupling undergo systematic changes in healthy aging and dementia (119–121). Taken together, these findings indicate that age related alterations in MEG power and temporal structure are embedded in large scale gradients of neuromodulation, cortical development and myelination, and cerebrovascular organization.

Several limitations should be considered when interpreting our results. First, our models are based on static, cross-sectional predictor maps and do not capture temporal dynamics at the level of neurotransmitter release, receptor trafficking, or gene expression, nor longitudinal changes in individuals. Second, our results rely on linear prediction and a step-wise feature selection procedure. Although this allowed us to identify a compact and interpretable feature set, true biological interactions are likely nonlinear, particularly across the lifespan (38,42,44,46), and the step wise approach favors complementary information, potentially down weighting features with redundant spatial distributions. Third, our predictor maps are derived from group-level PET, MRI, and transcriptomic datasets with heterogeneous sampling and measurement noise. Generalization of these atlases to diverse populations and across imaging sites necessitates further investigations (122). Finally, our work focuses on resting-state MEG. To what extent the same multiscale constraints shape task related dynamics and behavior will require future work. Despite these caveats, our findings provide a principled framework and hypotheses for future empirical and mechanistic studies. Usually, relationships between electrophysiology and neurochemical markers have been probed using MR spectroscopy, PET, pharmacological intervention, and pathological models, including benzodiazepine- and tiagabine-challenge studies and disease states that perturb excitation-inhibition balance. For example, the field has examined and controversially discussed links between certain, isolated neurotransmitter systems, prominently focusing on GABAergic and glutamatergic transmission and specific features of EEG/MEG signals in humans (123–131). Others have targeted cholinergic and dopaminergic effects on MEG/EEG (132) in pathological models (for example, (133–135)). These approaches are, however, constrained by the availability, specificity, and safety of perturbation and measurement tools for each receptor class. Thus, a comprehensive mapping across all major transmitter classes remains an important frontier for both basic science and clinical translation (136). Our study also complements recent efforts to build generative, biophysical models of regional variability in electrophysiological data (17) and behavioral performance of individual tasks (137).

In summary, our results demonstrate that specific markers sampling microstructure, transcriptomics, and neurotransmitter information accurately predicts parcel wise MEG power spectra and temporal autocorrelation across the cortex. We provide a comprehensive overview of predictive markers, their relative importance, and their spatial relationship with both spectral and temporal characteristics of electrophysiological signals. Moreover, we identify markers relevant for aging. Critically, this multivariate mapping can inform pharmacological and perturbational experiments by pinpointing neuromodulatory systems and structural motifs most likely to influence specific spectral and temporal neural signatures.

## Methods

### Sample and data source

This study used publicly available, cross-sectional data from the Cam-CAN initiative (51). Specifically, we examined healthy participants enrolled in the study’s second phase. The participant cohort included 350 individuals stratified by 50 participants per age decade, with equal numbers of males and females within each decade. Inclusion criteria required participants to be cognitively normal, without communication problems, free from major physical impairments, and without a history of neurological or psychiatric illness. Ethical oversight was provided by the Cambridgeshire 2 Research Ethics Committee, and the study adhered to principles outlined in the Declaration of Helsinki. Raw data are available through the Cam-CAN archive (https://camcan-archive.mrc-cbu.cam.ac.uk/dataaccess/).

### Structural MRI protocol

Anatomical images were obtained on a 3 T Siemens TIM Trio system equipped with a 32-channel head coil. T1-weighted images were acquired through 3D MPRAGE sequences with TR = 2,250 ms, TE = 2.99 ms, TI = 900 ms; FA = 9 deg; FOV = 256 × 240 × 192 mm; 1 mm isotropic; GRAPPA = 2; TA = 4 mins 32 s), and T2-weighted images from 3D SPACE sequences with TR = 2,800 ms,TE=408ms,TI=900ms;FOV=256×256×192mm;1mmisotropic; GRAPPA = 2; TA = 4 mins 30 s).

### MEG recording

Spontaneous brain activity was recorded using a whole-head MEG (Elekta Neuromag VectorView, Helsinki, Finland) with 306 channels (102 magnetometers and 204 planar gradiometers). The sampling frequency was set to 1 kHz, with an online high-pass filter of 0.03 Hz applied during acquisition. Recording sessions took place in a magnetically shielded chamber at a single institution (MRC Cognition and Brain Science Unit, Cambridge, UK). Each participant underwent an 8-minute 40-second resting-state session seated with eyes closed. Head position was tracked throughout the recording using four localization coils. Concurrent physiological monitoring included continuous electrocardiographic measurement (ECG) and horizontal and vertical electrooculogram (EOG).

### MEG/MRI processing

Cam-CAN provided initial processing of raw MEG data including temporal signal space separation (tSSS, Elekta Neuromag Maxfilter 2.2, 10 s window, 0.98 correlation limit). tSSS was applied for external artifact rejection, line noise removal (50 Hz and its harmonics), and correction for bad channels and head movements. Further preprocessing steps are described in detail elsewhere (38). In short, the data was resampled to 300 Hz, filtered (first-order Butterworth high-pass filter at 1 Hz), and segmented into 10-second trials using Fieldtrip (138). Following an automatic approach, we identified and rejected trials when containing movement artifacts (https://www.fieldtriptool-box.org/tutorial/automatic_artifact_rejection/). We low-pass filtered the data at 70 Hz (first-order Butterworth) and applied independent component analysis (ICA) to identify ocular and cardiac components. We rejected them based on their similarity to EOG signals (average coherence > 0.3 and amplitude correlation coefficient > 0.4) and coherence with the recorded ECG signal (average coherence > 0.3) or the averaged maximum peaks time-locked to the ECG (QRS complex). Before rejection, the selected components in each data set were manually checked. Because the ECG/EOG was noisy in a few cases, the relevant components had to be visually inspected and identified. All data underwent quality control and vigilance assessment according to the criteria of the American Academy of Sleep Medicine (https://aasm.org/). 30 clean and “awake” trials were then selected based on previous work that demonstrated good reliability for MEG spectral metrics (139). In the following, we only considered data from magnetometers (102 channels) since the content of both available channel types (magnetometers and gradiometers) has been highly similar after signal space separation (140).

### Individual source projection

Structural MR-images (T- and T2-weighted) were segmented, and surface extraction was applied using FreeSurfer (v6.0.0). We then projected MEG sensor data to the participant’s cortical surfaces based on the “fsaverage” template mesh provided by SUMA (141) and FreeSurfer. Here, each cortical surface was resampled to 1,002 vertices per brain hemisphere (ld factor = 10). Individual meshes were realigned to the Neuromag sensor space using anatomical landmarks that had been pre-identified during the Cam-CAN data acquisition protocol. Forward models characterizing the electromagnetic relationship between sources and sensors were generated employing a single-shell spherical approximation computed via Fieldtrip. We computed the covariance of processed MEG sensor data across trials and constructed spatial filters for each parcel of the Schaefer atlas (53) (regularization lambda = 5%, free-orientation forward solution). For each parcel, we concatenated the vertex filters, applied singular value decomposition, and retained the first singular vector (dominant source component) as the parcel time-series, resulting in 200 parcel-wise time-series per subject.

### Spectral power and temporal autocorrelation estimation

For each individual and parcel, power spectra were estimated in 1 Hz bins (1-60 Hz) using fast Fourier spectral analysis and multitapers (Discrete Prolate Spheroidal Sequences tapers) implemented in Fieltrip based on the cleaned, source-projected time-series (30 trials, 5 minutes of data). In addition, we computed temporal autocorrelation (AC) for up to 40 lags (∼3.3ms steps) using MATLAB’s ‘autocorr’ function following Box and colleagues (142). For each brain parcel, we obtained AC from time-series data via (inverse) Fast Fourier Transform, normalizing the function such that zero-lag autocorrelation equals one and remaining lags range between −1 and +1. As outlined in the following, we built average MEG maps for both metrics, power and AC, which were corrected with respect to age (linear and quadratic), sex, and estimated total intracranial volume (eTIV) using ordinary least squares. eTIV was obtained from FreeSurfer’s automated anatomical segmentation.

### Structural and molecular predictors

Predictors consisted of 55 regional maps (neurochemical and structural features) mapped to the Schaefer atlas (53) and concatenated into one predictor matrix. Maps summarizing cytoarchitectural properties were provided by the ENIGMA toolbox (27) and were originally computed on the BigBrain (https://bigbrain.loris.ca/main.php) across several cortical depths (Paquola et al. 2019). BigBrain is a microscopic resolution 3D brain model derived from a 65-year-old male (52) and techniques of Merker staining (143) for cell bodies. Statistical moments quantify different aspects of this histological data and its distribution, such as the amplitude reflecting the average intensity of the BigBrain staining profile (‘cyto-intensity’), irrespective of depth-wise differences, or the mean (‘cyto-mean’), summarizing the intensity across cortical depth. We also included the standard deviation (‘cyto-sd’), kurtosis (‘cyto-kurtosis’) and skewness (‘cyto-skewness’), which reflects laminar differentiation due to uneven cellular distribution (contrast between deep and superficial layers). The BigBrain gradient (‘cyto-gradient’) is built on histology-based microstructural covariance and indexes sensory-fugal neurostructural variation (66). We also derived a cortical hub measure from the ENIGMA toolbox based on the sum of all weighted cortico-cortical connections (degree centrality for each region, ‘struct-hub’), therefore reflecting structural network organization in the brain (27,144).

Further features were extracted using the neuromaps toolbox (26). Similar to previous work (28), available data at the surface level were transformed from their respective space to the fsLR32k space and parcellated into 200 regions based on the Schaefer 2018 atlas (53). This included maps characterizing microstructure, such as cortical thickness (‘thickness’) and cortical myelin (‘myelin’) estimated from T1w/T2w contrasts. We further included the first principal component of gene expression data derived from the Allen Human Brain Atlas (26,76). Also, we considered maps describing cortical organization along the sensorimotor-association axis (‘s-a-axis’) and expansion, evolutionary (‘ev-expansion’) and developmentally (‘dev-expansion’), as well as allometric scaling (‘cort-scaling’). We included maps probing brain metabolism such as cerebral bloodflow (‘cereb-bloodflow’), volume (‘cereb-volume’), oxygen (‘oxygen’), and glucose metabolism (‘glucose’).

Neurotransmitter receptor and transporter data were provided by neuromaps in the volumetric space (MNI152) and parcellated to the 200-parcel volumetric Schaefer atlas, since it is not recommended to transform PET volumes to the surface. We considered spatial maps for acetylcholine (‘α4β2’, ‘M1’, ‘VAChT’), cannabinoid (‘CB1’), dopamine (‘D1’, ‘D2’, ‘DAT’), glutamate (‘mGluR5’), and GABA (‘GABAa’, ‘GABAa_alpha5’), histamine (‘H3’), norepinephrine (‘NET’), opioid (‘MOR’, ‘KOR’), serotonin (‘5-HT1a’, ‘5-HT1b’, ‘5-HT2a’, ‘5-HT4’, ‘5-HT6’, ‘5-HTT’). Moreover, a map quantifying synapse density was included (tracer binding to the synaptic vesicle glycoprotein 2A (‘SV2A’)), a translocator protein (‘TSPO’), as well as enzymes such as (‘COX-1’) and histone deacetylase (‘HDAC’). References for the respective maps are listed in **Table 1**.

### Prediction of parcel-wise power spectra

We used partial least squares regression (PLSR) to predict the parcel-wise average power spectrum (1-60 Hz) from the set of structural and functional brain maps (n=55). Before prediction, we normalized the raw power spectrum of each parcel by subtracting its global mean (across frequencies) and applied a log-transformation. This removes the broadband offset for each parcel and centers the spectra around zero. For an initial assessment of explained variance, we ran PLSR on the full dataset with 30 components (latent variables). We used 10 components throughout the subsequent analyses, as this provided the optimal trade-off between variance explained and number of components. We then implemented 10-fold cross-validation and 50 repetitions for robustness. To avoid information leakage, predictors were standardized inside the cross-validation loop. Specifically, within each fold, the predictor maps were standardized using the mean and standard deviation of the training data (180 parcels per map), and the held-out data (20 parcels per map) was transformed using those training statistics. Moreover, within each fold, we applied z-scoring to each frequency bin computed across parcels in the training set. This procedure removes the global similarity of the spectrum across parcels, hence enhancing differences between parcels.

### Prediction of parcel-wise autocorrelation function

We applied a similar procedure to assess temporal autocorrelation (AC) in our MEG data, considering the autocorrelation function over up to 40 time-lags (corresponding to shifts of ∼3,3 ms). Hence, we used PLSR to predict parcel-wise AC (1-40 lags) from the same set of structural and functional brain maps (n=55) and kept the same number of components (n=10) as for power to ensure comparability. We again implemented 10-fold cross-validation and 50 repetitions, and predictors were standardized inside the cross-validation loop as described above. Inside each fold, we subtracted the AC mean across training parcels within each time-lag to remove global similarity across parcels. Therefore, in the end, parcel-wise relative deviations are predicted, not the raw autocorrelation function.

### Evaluation of model performance

Model performance was summarized using: (i) overall prediction R² computed as 1 - normalized mean standard error (or the proportional improvement of the model over simply predicting the mean, (145)) and (ii) rank accuracy obtained by computing the parcel-by-parcel correlation between observed and predicted spectra and taking the median of tied ranks across parcels. In essence, this yields the median identifiability of individual parcels. A rank of 200 would correspond to a perfect match between the predicted and real spectrum or function of a parcel. To assess significance, we generated null models using spin-tests (93) that spherically rotate and reassign values to brain parcels. For each spin (n = 1000), we repeated the respective PLS approach and tested the empirical outcome against the generated null distribution. To assess the importance of single predictors or biological maps, we computed variable importance in projection (VIP) scores per fold using the PLS weights and latent scores and averaged VIPs across folds and repetitions to obtain a stable ranking of predictors. A VIP score > 1 indicates a strong contribution of a predictor to the PLS model.

### Incremental prediction

Instead of using all predictors at once in a multivariate fashion, we have tested in the next step how many of them and in which combination predict MEG power or autocorrelation achieve the same accuracy as compared to the full model. We therefore implemented an approach by adding predictive maps incrementally to the PLSR model. Hence, the map yielding the highest prediction R² will be identified first, and the next map will be added based on its combination, adding up to the next highest R². To keep consistency, we increased the number of PLS components gradually after every fifth predictor.

### Age-specific predictions / Individual deviations and aging effects

Next, we aimed at identifying biological correlates of individual aging effects in the MEG data. Given the balanced age distribution of the available MEG sample (18-88 years, n=50 individuals per decade), we computed individual z-scored deviations in the power spectrum or autocorrelation function from the average maps. These deviations represent age-related effects across the adult lifespan and can be used for inferring biological underpinnings. Hence, we set up individual PLSR models with individual MEG deviations (200 parcels by 1-60 Hz bins and 1-40 time lags, respectively) as input and the full set of predictor maps (200 parcels by 55 maps). For comparison, we kept the number of PLS components (n=10) constant and derived subject-level model performance and beta weights. To understand which predictors align with these aging effects best, we then correlated individual beta weights for each map with chronological age using Pearson correlation. Hence, the magnitude of the correlation coefficient indicates the relationship between predictor maps and age effects in the MEG data.

## Acknowledgements

Data collection and sharing for this project was provided by the Cambridge Center for Ageing and Neuroscience (CamCAN). Cam-CAN funding was provided by the UK Biotechnology and Biological Sciences Research Council (Grant number BB/H008217/1), together with support from the UK Medical Research Council and the University of Cambridge, UK. This work was further supported by the German Research Foundation (DFG; project grant no 491157081 to J.G.; SFB/TRR 393 project grant no 521379614 to J.G. and U.D.; grant FOR2107 DA1151/5-1, DA1151/5-2, DA1151/9-1, DA1151/10-1, DA1151/11-1 to UD); and the Interdisciplinary Center for Clinical Research (IZKF) of the medical faculty of Münster (grant Dan3/016/26).

## Author contributions

Conceptualization: CS, JG

Methodology: CS, JG

Investigation: CS, JG

Visualization: CS

Supervision: JG

Writing—original draft: CS, JG

Writing—review & editing: CS, JG, UD

## Data and materials availability

Anonymized raw data were provided by the Cambridge Center for Ageing and Neuroscience and available at Cam-CAN Data Portal (https://camcan-archive.mrc-cbu.cam.ac.uk/dataaccess/). Structural and anatomical maps are openly available and can be derived via https://enigma-toolbox.readthedocs.io/en/latest/index.html and https://netneurolab.github.io/neuromaps/index.html. Analysis code and results are accessible at https://github.com/chstier/StructFunc.git

## References

1. Suárez LE, Markello RD, Betzel RF, Misic B. Linking structure and function in macroscale brain networks. Trends Cogn Sci (Regul Ed). 2020 Apr;24(4):302–15.

2. van den Heuvel MP, Mandl RCW, Kahn RS, Hulshoff Pol HE. Functionally linked resting-state networks reflect the underlying structural connectivity architecture of the human brain. Hum Brain Mapp. 2009 Oct;30(10):3127–41.

3. Mišić B, Betzel RF, de Reus MA, van den Heuvel MP, Berman MG, McIntosh AR, et al. Network-Level Structure-Function Relationships in Human Neocortex. Cereb Cortex. 2016 Jul;26(7):3285–96.

4. Messé A, Rudrauf D, Benali H, Marrelec G. Relating structure and function in the human brain: relative contributions of anatomy, stationary dynamics, and non-stationarities. PLoS Comput Biol. 2014 Mar 20;10(3):e1003530.

5. Fukushima M, Betzel RF, He Y, van den Heuvel MP, Zuo X-N, Sporns O. Structure-function relationships during segregated and integrated network states of human brain functional connectivity. Brain Struct Funct. 2018 Apr;223(3):1091–106.

6. Yang Y, Zheng Z, Liu L, Zheng H, Zhen Y, Zheng Y, et al. Enhanced brain structure-function tethering in transmodal cortex revealed by high-frequency eigenmodes. Nat Commun. 2023 Oct 24;14(1):6744.

7. Preti MG, Van De Ville D. Decoupling of brain function from structure reveals regional behavioral specialization in humans. Nat Commun. 2019 Oct 18;10(1):4747.

8. Baum GL, Cui Z, Roalf DR, Ciric R, Betzel RF, Larsen B, et al. Development of structure-function coupling in human brain networks during youth. Proc Natl Acad Sci USA. 2020 Jan 7;117(1):771–8.

9. Vázquez-Rodríguez B, Suárez LE, Markello RD, Shafiei G, Paquola C, Hagmann P, et al. Gradients of structure-function tethering across neocortex. Proc Natl Acad Sci USA. 2019 Oct 15;116(42):21219–27.

10. Wu D, Fan L, Song M, Wang H, Chu C, Yu S, et al. Hierarchy of Connectivity-Function Relationship of the Human Cortex Revealed through Predicting Activity across Functional Domains. Cereb Cortex. 2020 Jun 30;30(8):4607–16.

11. Huntenburg JM, Bazin P-L, Goulas A, Tardif CL, Villringer A, Margulies DS. A systematic relationship between functional connectivity and intracortical myelin in the human cerebral cortex. Cereb Cortex. 2017 Feb 1;27(2):981–97.

12. Yang Y, Tang S, Wang X, Zhen Y, Zheng Y, Zheng H, et al. Eigenmode-based approach reveals a decline in brain structure-function liberality across the human lifespan. Commun Biol. 2023 Nov 7;6(1):1128.

13. Liu Z-Q, Vázquez-Rodríguez B, Spreng RN, Bernhardt BC, Betzel RF, Misic B. Time-resolved structure-function coupling in brain networks. Commun Biol. 2022 Jun 2;5(1):532.

14. Buzsáki G, Anastassiou CA, Koch C. The origin of extracellular fields and currents--EEG, ECoG, LFP and spikes. Nat Rev Neurosci. 2012 May 18;13(6):407–20.

15. Smit DJA, Wright MJ, Meyers JL, Martin NG, Ho YYW, Malone SM, et al. Genome-wide association analysis links multiple psychiatric liability genes to oscillatory brain activity. Hum Brain Mapp. 2018 Nov;39(11):4183–95.

16. Smit DJA, Posthuma D, Boomsma DI, Geus EJC. Heritability of background EEG across the power spectrum. Psychophysiology. 2005 Nov;42(6):691–7.

17. Stoof UM, Friston KJ, Tisdall M, Cooray GK, Rosch RE. Topographic variation in human neurotransmitter receptor densities explains differences in intracranial EEG spectra. Hum Brain Mapp. 2025 Nov;46(16):e70393.

18. Keitel A, Gross J. Individual Human Brain Areas Can Be Identified from Their Characteristic Spectral Activation Fingerprints. PLoS Biol. 2016 Jun 29;14(6):e1002498.

19. Sydnor VJ, Larsen B, Bassett DS, Alexander-Bloch A, Fair DA, Liston C, et al. Neurodevelopment of the association cortices: Patterns, mechanisms, and implications for psychopathology. Neuron. 2021 Sep 15;109(18):2820–46.

20. Huntenburg JM, Bazin P-L, Margulies DS. Large-Scale Gradients in Human Cortical Organization. Trends Cogn Sci (Regul Ed). 2018 Jan;22(1):21–31.

21. Gao R, van den Brink RL, Pfeffer T, Voytek B. Neuronal timescales are functionally dynamic and shaped by cortical microarchitecture. eLife. 2020 Nov 23;9.

22. Fehring J, Balestrieri E, Focke N, Dannlowski U, Stier C, Gross J. Neurophysiological correlates of cortical hierarchy across the lifespan. BioRxiv. 2024 Jul 17;

23. Mahjoory K, Schoffelen J-M, Keitel A, Gross J. The frequency gradient of human resting-state brain oscillations follows cortical hierarchies. eLife. 2020 Aug 21;9.

24. Hunt BAE, Tewarie PK, Mougin OE, Geades N, Jones DK, Singh KD, et al. Relationships between cortical myeloarchitecture and electrophysiological networks. Proc Natl Acad Sci USA. 2016 Nov 22;113(47):13510–5.

25. Karahan E, Tait L, Si R, Özkan A, Szul MJ, Graham KS, et al. The interindividual variability of multimodal brain connectivity maintains spatial heterogeneity and relates to tissue microstructure. Commun Biol. 2022 Sep 23;5(1):1007.

26. Markello RD, Hansen JY, Liu Z-Q, Bazinet V, Shafiei G, Suárez LE, et al. neuromaps: structural and functional interpretation of brain maps. Nat Methods. 2022 Nov;19(11):1472–9.

27. Larivière S, Paquola C, Park B-Y, Royer J, Wang Y, Benkarim O, et al. The ENIGMA Toolbox: multiscale neural contextualization of multisite neuroimaging datasets. Nat Methods. 2021 Jul;18(7):698–700.

28. Shafiei G, Fulcher BD, Voytek B, Satterthwaite TD, Baillet S, Misic B. Neurophysiological signatures of cortical micro-architecture. BioRxiv. 2023 Jan 23;

29. Hansen JY, Shafiei G, Markello RD, Smart K, Cox SML, Nørgaard M, et al. Mapping neurotransmitter systems to the structural and functional organization of the human neocortex. Nat Neurosci. 2022 Nov;25(11):1569–81.

30. Siebenhühner F, Palva JM, Palva S. Linking the microarchitecture of neurotransmitter systems to large-scale MEG resting state networks. iScience. 2024 Oct;111111.

31. Park B-Y, Paquola C, Bethlehem RAI, Benkarim O, Neuroscience in Psychiatry Network (NSPN) Consortium, Mišić B, et al. Adolescent development of multiscale structural wiring and functional interactions in the human connectome. Proc Natl Acad Sci USA. 2022 Jul 5;119(27):e2116673119.

32. Zamani Esfahlani F, Faskowitz J, Slack J, Mišić B, Betzel RF. Local structure-function relationships in human brain networks across the lifespan. Nat Commun. 2022 Apr 19;13(1):2053.

33. Pedersen R, Geerligs L, Andersson M, Gorbach T, Avelar-Pereira B, Wåhlin A, et al. When functional blurring becomes deleterious: Reduced system segregation is associated with less white matter integrity and cognitive decline in aging. Neuroimage. 2021 Nov 15;242:118449.

34. Jauny G, Mijalkov M, Canal-Garcia A, Volpe G, Pereira J, Eustache F, et al. Linking structural and functional changes during aging using multilayer brain network analysis. Commun Biol. 2024 Feb 28;7(1):239.

35. Hinault T, Mijalkov M, Pereira JB, Volpe G, Bakke A, Courtney SM. Age-related differences in network structure and dynamic synchrony of cognitive control. Neuroimage. 2021 Aug 1;236:118070.

36. Hinault T, Kraut M, Bakker A, Dagher A, Courtney SM. Disrupted Neural Synchrony Mediates the Relationship between White Matter Integrity and Cognitive Performance in Older Adults. Cereb Cortex. 2020 Sep 3;30(10):5570–82.

37. Kalpouzos G, Persson J. Structure-function relationships in the human aging brain: An account of cross-sectional and longitudinal multimodal neuroimaging studies. Cortex. 2025 Feb;183:274–89.

38. Stier C, Braun C, Focke NK. Adult lifespan trajectories of neuromagnetic signals and interrelations with cortical thickness. Neuroimage. 2023 Sep;278:120275.

39. Barry RJ, De Blasio FM. EEG differences between eyes-closed and eyes-open resting remain in healthy ageing. Biol Psychol. 2017 Oct;129:293–304.

40. Coquelet N, Wens V, Mary A, Niesen M, Puttaert D, Ranzini M, et al. Changes in electrophysiological static and dynamic human brain functional architecture from childhood to late adulthood. Sci Rep. 2020 Nov 4;10(1):18986.

41. Duffy FH, Albert MS, McAnulty G, Garvey AJ. Agelrelated differences in brain electrical activity of healthy subjects. Annals of Neurology: Official Journal of the American Neurological Association and the Child Neurology Society. 1984;16(4):430–8.

42. Gómez C, Pérez-Macías JM, Poza J, Fernández A, Hornero R. Spectral changes in spontaneous MEG activity across the lifespan. J Neural Eng. 2013 Dec;10(6):066006.

43. Ustinin M, Boyko A, Rykunov S. Healthy aging changes in conventional frequency bands of neuroelectric brain activity reconstructed from resting-state MEG. Geroscience. 2025 Jun;47(3):4093–108.

44. Doval S, López-Sanz D, Bruña R, Cuesta P, Antón-Toro L, Taguas I, et al. When maturation is not linear: brain oscillatory activity in the process of aging as measured by electrophysiology. Brain Topogr. 2024 Nov;37(6):1068–88.

45. Hunt BAE, Wong SM, Vandewouw MM, Brookes MJ, Dunkley BT, Taylor MJ. Spatial and spectral trajectories in typical neurodevelopment from childhood to middle age. Netw Neurosci. 2019 Mar 1;3(2):497–520.

46. Rempe MP, Ott LR, Picci G, Penhale SH, Christopher-Hayes NJ, Lew BJ, et al. Spontaneous cortical dynamics from the first years to the golden years. Proc Natl Acad Sci USA. 2023 Jan 24;120(4):e2212776120.

47. Perinelli A, Assecondi S, Tagliabue CF, Mazza V. Power shift and connectivity changes in healthy aging during resting-state EEG. Neuroimage. 2022 Aug 1;256:119247.

48. Shinn M, Hu A, Turner L, Noble S, Preller KH, Ji JL, et al. Functional brain networks reflect spatial and temporal autocorrelation. Nat Neurosci. 2023 May;26(5):867–78.

49. Stier C, Balestrieri E, Fehring J, Focke NK, Wollbrink A, Dannlowski U, et al. Temporal autocorrelation is predictive of age-An extensive MEG time-series analysis. Proc Natl Acad Sci USA. 2025 Feb 25;122(8):e2411098122.

50. Taylor JR, Williams N, Cusack R, Auer T, Shafto MA, Dixon M, et al. The Cambridge Centre for Ageing and Neuroscience (Cam-CAN) data repository: Structural and functional MRI, MEG, and cognitive data from a cross-sectional adult lifespan sample. Neuroimage. 2017 Jan;144(Pt B):262–9.

51. Shafto MA, Tyler LK, Dixon M, Taylor JR, Rowe JB, Cusack R, et al. The Cambridge Centre for Ageing and Neuroscience (Cam-CAN) study protocol: a cross-sectional, lifespan, multidisciplinary examination of healthy cognitive ageing. BMC Neurol. 2014;14.

52. Amunts K, Lepage C, Borgeat L, Mohlberg H, Dickscheid T, Rousseau M-É, et al. BigBrain: an ultrahigh-resolution 3D human brain model. Science. 2013 Jun 21;340(6139):1472–5.

53. Schaefer A, Kong R, Gordon EM, Laumann TO, Zuo X-N, Holmes AJ, et al. Local-Global Parcellation of the Human Cerebral Cortex from Intrinsic Functional Connectivity MRI. Cereb Cortex. 2018 Sep 1;28(9):3095–114.

54. Savli M, Bauer A, Mitterhauser M, Ding Y-S, Hahn A, Kroll T, et al. Normative database of the serotonergic system in healthy subjects using multi-tracer PET. Neuroimage. 2012 Oct 15;63(1):447–59.

55. Gallezot J-D, Nabulsi N, Neumeister A, Planeta-Wilson B, Williams WA, Singhal T, et al. Kinetic modeling of the serotonin 5-HT(1B) receptor radioligand [(11)C]P943 in humans. J Cereb Blood Flow Metab. 2010 Jan;30(1):196–210.

56. Beliveau V, Ganz M, Feng L, Ozenne B, Højgaard L, Fisher PM, et al. A High-Resolution In Vivo Atlas of the Human Brain’s Serotonin System. J Neurosci. 2017 Jan 4;37(1):120–8.

57. Radhakrishnan R, Nabulsi N, Gaiser E, Gallezot J-D, Henry S, Planeta B, et al. Age-Related Change in 5-HT6 Receptor Availability in Healthy Male Volunteers Measured with 11C-GSK215083 PET. J Nucl Med. 2018 Sep;59(9):1445–50.

58. Fazio P, Schain M, Varnäs K, Halldin C, Farde L, Varrone A. Mapping the distribution of serotonin transporter in the human brainstem with high-resolution PET: Validation using postmortem autoradiography data. Neuroimage. 2016 Jun;133:313–20.

59. Hillmer AT, Esterlis I, Gallezot JD, Bois F, Zheng MQ, Nabulsi N, et al. Imaging of cerebral α4β2* nicotinic acetylcholine receptors with (-)-[(18)F]Flubatine PET: Implementation of bolus plus constant infusion and sensitivity to acetylcholine in human brain. Neuroimage. 2016 Nov 1;141:71–80.

60. Normandin MD, Zheng M-Q, Lin K-S, Mason NS, Lin S-F, Ropchan J, et al. Imaging the cannabinoid CB1 receptor in humans with [11C]OMAR: assessment of kinetic analysis methods, test-retest reproducibility, and gender differences. J Cereb Blood Flow Metab. 2015 Aug;35(8):1313–22.

61. Vaishnavi SN, Vlassenko AG, Rundle MM, Snyder AZ, Mintun MA, Raichle ME. Regional aerobic glycolysis in the human brain. Proc Natl Acad Sci USA. 2010 Oct 12;107(41):17757–62.

62. Satterthwaite TD, Shinohara RT, Wolf DH, Hopson RD, Elliott MA, Vandekar SN, et al. Impact of puberty on the evolution of cerebral perfusion during adolescence. Proc Natl Acad Sci USA. 2014 Jun 10;111(23):8643–8.

63. Reardon PK, Seidlitz J, Vandekar S, Liu S, Patel R, Park MTM, et al. Normative brain size variation and brain shape diversity in humans. Science. 2018 Jun 15;360(6394):1222–7.

64. Greve D, Ganz M, Nørgaard M, Matheson G, Pernet C, Galassi A, et al. PS13 Molecular Imaging Brain Atlas (Kim 2021). Openneuro. 2023;

65. Kim M-J, Lee J-H, Juarez Anaya F, Hong J, Miller W, Telu S, et al. First-in-human evaluation of [11C]PS13, a novel PET radioligand, to quantify cyclooxygenase-1 in the brain. Eur J Nucl Med Mol Imaging. 2020 Dec;47(13):3143–51.

66. Paquola C, Vos De Wael R, Wagstyl K, Bethlehem RAI, Hong S-J, Seidlitz J, et al. Microstructural and functional gradients are increasingly dissociated in transmodal cortices. PLoS Biol. 2019 May 20;17(5):e3000284.

67. Kaller S, Rullmann M, Patt M, Becker G-A, Luthardt J, Girbardt J, et al. Test-retest measurements of dopamine D1-type receptors using simultaneous PET/MRI imaging. Eur J Nucl Med Mol Imaging. 2017 Jun;44(6):1025–32.

68. Jaworska N, Cox SML, Tippler M, Castellanos-Ryan N, Benkelfat C, Parent S, et al. Extra-striatal D2/3 receptor availability in youth at risk for addiction. Neuropsychopharmacology. 2020 Aug;45(9):1498–505.

69. Malén T, Karjalainen T, Isojärvi J, Vehtari A, Bürkner P-C, Putkinen V, et al. Atlas of type 2 dopamine receptors in the human brain: Age and sex dependent variability in a large PET cohort. Neuroimage. 2022 Jul 15;255:119149.

70. Smith CT, Crawford JL, Dang LC, Seaman KL, San Juan MD, Vijay A, et al. Partial-volume correction increases estimated dopamine D2-like receptor binding potential and reduces adult age differences. J Cereb Blood Flow Metab. 2019 May;39(5):822–33.

71. Dukart J, Holiga Š, Chatham C, Hawkins P, Forsyth A, McMillan R, et al. Cerebral blood flow predicts differential neurotransmitter activity. Sci Rep. 2018 Mar 6;8(1):4074.

72. Sasaki T, Ito H, Kimura Y, Arakawa R, Takano H, Seki C, et al. Quantification of dopamine transporter in human brain using PET with 18F-FE-PE2I. J Nucl Med. 2012 Jul;53(7):1065–73.

73. Hill J, Inder T, Neil J, Dierker D, Harwell J, Van Essen D. Similar patterns of cortical expansion during human development and evolution. Proc Natl Acad Sci USA. 2010 Jul 20;107(29):13135–40.

74. Lukow PB, Martins D, Veronese M, Vernon AC, McGuire P, Turkheimer FE, et al. Cellular and molecular signatures of in vivo imaging measures of GABAergic neurotransmission in the human brain. Commun Biol. 2022 Apr 19;5(1):372.

75. Nørgaard M, Beliveau V, Ganz M, Svarer C, Pinborg LH, Keller SH, et al. A high-resolution in vivo atlas of the human brain’s benzodiazepine binding site of GABAA receptors. Neuroimage. 2021 May 15;232:117878.

76. Hawrylycz MJ, Lein ES, Guillozet-Bongaarts AL, Shen EH, Ng L, Miller JA, et al. An anatomically comprehensive atlas of the adult human brain transcriptome. Nature. 2012 Sep 20;489(7416):391–9.

77. Markello RD, Arnatkeviciute A, Poline J-B, Fulcher BD, Fornito A, Misic B. Standardizing workflows in imaging transcriptomics with the abagen toolbox. eLife. 2021 Nov 16;10.

78. Gallezot J-D, Planeta B, Nabulsi N, Palumbo D, Li X, Liu J, et al. Determination of receptor occupancy in the presence of mass dose: [11C]GSK189254 PET imaging of histamine H3 receptor occupancy by PF-03654746. J Cereb Blood Flow Metab. 2017 Mar;37(3):1095–107.

79. Wey H-Y, Gilbert TM, Zürcher NR, She A, Bhanot A, Taillon BD, et al. Insights into neuroepigenetics through human histone deacetylase PET imaging. Sci Transl Med. 2016 Aug 10;8(351):351ra106.

80. Vijay A, Cavallo D, Goldberg A, de Laat B, Nabulsi N, Huang Y, et al. PET imaging reveals lower kappa opioid receptor availability in alcoholics but no effect of age. Neuropsychopharmacology. 2018 Dec;43(13):2539–47.

81. Naganawa M, Nabulsi N, Henry S, Matuskey D, Lin S-F, Slieker L, et al. First-in-Human Assessment of 11C-LSN3172176, an M1 Muscarinic Acetylcholine Receptor PET Radiotracer. J Nucl Med. 2021 Apr;62(4):553–60.

82. DuBois JM, Rousset OG, Rowley J, Porras-Betancourt M, Reader AJ, Labbe A, et al. Characterization of age/sex and the regional distribution of mGluR5 availability in the healthy human brain measured by high-resolution [(11)C]ABP688 PET. Eur J Nucl Med Mol Imaging. 2016 Jan;43(1):152–62.

83. Smart K, Cox SML, Scala SG, Tippler M, Jaworska N, Boivin M, et al. Sex differences in [11C]ABP688 binding: a positron emission tomography study of mGlu5 receptors. Eur J Nucl Med Mol Imaging. 2019 May;46(5):1179–83.

84. Kantonen T, Karjalainen T, Isojärvi J, Nuutila P, Tuisku J, Rinne J, et al. Interindividual variability and lateralization of μ-opioid receptors in the human brain. Neuroimage. 2020 Aug 15;217:116922.

85. Glasser MF, Coalson TS, Robinson EC, Hacker CD, Harwell J, Yacoub E, et al. A multi-modal parcellation of human cerebral cortex. Nature. 2016 Aug 11;536(7615):171–8.

86. Ding Y-S, Singhal T, Planeta-Wilson B, Gallezot J-D, Nabulsi N, Labaree D, et al. PET imaging of the effects of age and cocaine on the norepinephrine transporter in the human brain using (S,S)-[(11)C]O-methylreboxetine and HRRT. Synapse. 2010 Jan;64(1):30–8.

87. Van Essen DC, Smith SM, Barch DM, Behrens TEJ, Yacoub E, Ugurbil K, et al. The WU-Minn Human Connectome Project: an overview. Neuroimage. 2013 Oct 15;80:62–79.

88. Finnema SJ, Nabulsi NB, Mercier J, Lin S-F, Chen M-K, Matuskey D, et al. Kinetic evaluation and test-retest reproducibility of [11C]UCB-J, a novel radioligand for positron emission tomography imaging of synaptic vesicle glycoprotein 2A in humans. J Cereb Blood Flow Metab. 2018 Nov;38(11):2041–52.

89. Naganawa M, Li S, Nabulsi N, Henry S, Zheng M-Q, Pracitto R, et al. First-in-Human Evaluation of 18F-SynVesT-1, a Radioligand for PET Imaging of Synaptic Vesicle Glycoprotein 2A. J Nucl Med. 2021 Apr;62(4):561–7.

90. Lois C, González I, Izquierdo-García D, Zürcher NR, Wilkens P, Loggia ML, et al. Neuroinflammation in Huntington’s Disease: New Insights with 11C-PBR28 PET/MRI. ACS Chem Neurosci. 2018 Nov 21;9(11):2563–71.

91. Gonzalez I, Lois C. hookerlab/huntington-with-pbr28: ACS Chemical Neuroscience submission. Zenodo. 2018;

92. Aghourian M, Legault-Denis C, Soucy JP, Rosa-Neto P, Gauthier S, Kostikov A, et al. Quantification of brain cholinergic denervation in Alzheimer’s disease using PET imaging with [18F]-FEOBV. Mol Psychiatry. 2017 Nov;22(11):1531–8.

93. Alexander-Bloch AF, Shou H, Liu S, Satterthwaite TD, Glahn DC, Shinohara RT, et al. On testing for spatial correspondence between maps of human brain structure and function. Neuroimage. 2018 Sep;178:540–51.

94. Lundqvist M, Bastos AM, Miller EK. Preservation and Changes in Oscillatory Dynamics across the Cortical Hierarchy. J Cogn Neurosci. 2020 Oct;32(10):2024–35.

95. Cheyne D, Bells S, Ferrari P, Gaetz W, Bostan AC. Self-paced movements induce high-frequency gamma oscillations in primary motor cortex. Neuroimage. 2008 Aug 1;42(1):332–42.

96. Capilla A, Schoffelen J-M, Paterson G, Thut G, Gross J. Dissociated α-band modulations in the dorsal and ventral visual pathways in visuospatial attention and perception. Cereb Cortex. 2014 Feb;24(2):550–61.

97. Michalareas G, Vezoli J, van Pelt S, Schoffelen J-M, Kennedy H, Fries P. Alpha-Beta and Gamma Rhythms Subserve Feedback and Feedforward Influences among Human Visual Cortical Areas. Neuron. 2016 Jan 20;89(2):384–97.

98. Ploner M, Sorg C, Gross J. Brain rhythms of pain. Trends Cogn Sci (Regul Ed). 2017 Feb;21(2):100–10.

99. Burt JB, Demirtaş M, Eckner WJ, Navejar NM, Ji JL, Martin WJ, et al. Hierarchy of transcriptomic specialization across human cortex captured by structural neuroimaging topography. Nat Neurosci. 2018 Sep;21(9):1251–9.

100. Shafiei G, Keller AS, Bertolero M, Shanmugan S, Bassett DS, Chen AA, et al. Generalizable links between symptoms of borderline personality disorder and functional connectivity. BioRxiv. 2023 Aug 6;

101. Cole RH, Allichon M-C, Joffe ME. Opioid Receptors Modulate Inhibition within the Prefrontal Cortex through Dissociable Cellular and Molecular Mechanisms. J Neurosci. 2025 Jul 2;45(27).

102. Smith SJ, Sümbül U, Graybuck LT, Collman F, Seshamani S, Gala R, et al. Single-cell transcriptomic evidence for dense intracortical neuropeptide networks. eLife. 2019 Nov 11;8.

103. Khintchine A. Korrelationstheorie der stationlren stochastischen Prozesse. Math Ann. 1934 Dec;109(1):604–15.

104. Balestrieri E, Chalas N, Stier C, Fehring J, Gil Ávila C, Dannlowski U, et al. Beyond oscillations—Toward a richer characterization of brain states. Imaging Neuroscience. 2025 Feb 27;3.

105. Phillis JW, Horrocks LA, Farooqui AA. Cyclooxygenases, lipoxygenases, and epoxygenases in CNS: their role and involvement in neurological disorders. Brain Res Rev. 2006 Sep;52(2):201–43.

106. Shukuri M, Mawatari A, Ohno M, Suzuki M, Doi H, Watanabe Y, et al. Detection of Cyclooxygenase-1 in Activated Microglia During Amyloid Plaque Progression: PET Studies in Alzheimer’s Disease Model Mice. J Nucl Med. 2016 Feb;57(2):291–6.

107. Bazan NG, Colangelo V, Lukiw WJ. Prostaglandins and other lipid mediators in Alzheimer’s disease. Prostaglandins Other Lipid Mediat. 2002 Aug;68–69:197–210.

108. Liu Y, Traba JE, Lüscher C. Striatal crosstalk between dopamine and serotonin systems. 2025 Jul 16;

109. Gasiorowska A, Wydrych M, Drapich P, Zadrozny M, Steczkowska M, Niewiadomski W, et al. The biology and pathobiology of glutamatergic, cholinergic, and dopaminergic signaling in the aging brain. Front Aging Neurosci. 2021 Jul 13;13:654931.

110. Bartus RT, Dean RL, Beer B, Lippa AS. The cholinergic hypothesis of geriatric memory dysfunction. Science. 1982 Jul 30;217(4558):408–14.

111. Gedankien T, Tan RJ, Qasim SE, Moore H, McDonagh D, Jacobs J, et al. Acetylcholine modulates the temporal dynamics of human theta oscillations during memory. Nat Commun. 2023 Aug 30;14(1):5283.

112. Whitehouse PJ, Price DL, Clark AW, Coyle JT, DeLong MR. Alzheimer disease: evidence for selective loss of cholinergic neurons in the nucleus basalis. Ann Neurol. 1981 Aug;10(2):122–6.

113. Arendt T, Bigl V, Arendt A, Tennstedt A. Loss of neurons in the nucleus basalis of Meynert in Alzheimer’s disease, paralysis agitans and Korsakoff’s Disease. Acta Neuropathol. 1983;61(2):101–8.

114. Whitehouse PJ, Hedreen JC, White CL, Price DL. Basal forebrain neurons in the dementia of Parkinson disease. Ann Neurol. 1983 Mar;13(3):243–8.

115. Kasper J, Caspers S, Lotter LD, Hoffstaedter F, Eickhoff SB, Dukart J. Resting-State Changes in Aging and Parkinson’s Disease Are Shaped by Underlying Neurotransmission: A Normative Modeling Study. Biol Psychiatry Cogn Neurosci Neuroimaging. 2024 Oct;9(10):986–97.

116. Grydeland H, Vértes PE, Váša F, Romero-Garcia R, Whitaker K, Alexander-Bloch AF, et al. Waves of Maturation and Senescence in Micro-structural MRI Markers of Human Cortical Myelination over the Lifespan. Cereb Cortex. 2019 Mar 1;29(3):1369–81.

117. de Faria O, Pivonkova H, Varga B, Timmler S, Evans KA, Káradóttir RT. Periods of synchronized myelin changes shape brain function and plasticity. Nat Neurosci. 2021 Nov;24(11):1508–21.

118. Marek S, Tervo-Clemmens B, Klein N, Foran W, Ghuman AS, Luna B. Adolescent development of cortical oscillations: Power, phase, and support of cognitive maturation. PLoS Biol. 2018 Nov 30;16(11):e2004188.

119. Farahani A, Liu Z-Q, Ceballos EG, Hansen JY, Wennberg K, Zeighami Y, et al. Mapping cerebral blood perfusion and its links to multi-scale brain organization across the human lifespan. PLoS Biol. 2025 Jul 29;23(7):e3003277.

120. Tarantini S, Tran CHT, Gordon GR, Ungvari Z, Csiszar A. Impaired neurovascular coupling in aging and Alzheimer’s disease: Contribution of astrocyte dysfunction and endothelial impairment to cognitive decline. Exp Gerontol. 2017 Aug;94:52–8.

121. Zhang N, Gordon ML, Goldberg TE. Cerebral blood flow measured by arterial spin labeling MRI at resting state in normal aging and Alzheimer’s disease. Neurosci Biobehav Rev. 2017 Jan;72:168–75.

122. Hansen JY, Tuisku J, Johansson J, Chang Z, McGinnity CJ, Beliveau V, et al. Inter-individual variability of neurotransmitter receptor and transporter density in the human brain. Brain Struct Funct. 2026 Jan 13;231(1):13.

123. Baumgarten TJ, Oeltzschner G, Hoogenboom N, Wittsack H-J, Schnitzler A, Lange J. Beta Peak Frequencies at Rest Correlate with Endogenous GABA+/Cr Concentrations in Sensorimotor Cortex Areas. PLoS ONE. 2016 Jun 3;11(6):e0156829.

124. Muthukumaraswamy SD, Edden RAE, Jones DK, Swettenham JB, Singh KD. Resting GABA concentration predicts peak gamma frequency and fMRI amplitude in response to visual stimulation in humans. Proc Natl Acad Sci USA. 2009 May 19;106(20):8356–61.

125. Kujala J, Jung J, Bouvard S, Lecaignard F, Lothe A, Bouet R, et al. Gamma oscillations in V1 are correlated with GABA(A) receptor density: A multi-modal MEG and Flumazenil-PET study. Sci Rep. 2015 Nov 17;5:16347.

126. Kujala J, Ciumas C, Jung J, Bouvard S, Lecaignard F, Lothe A, et al. GABAergic inhibition shapes behavior and neural dynamics in human visual working memory. Cereb Cortex. 2024 Jan 31;34(2).

127. Cousijn H, Haegens S, Wallis G, Near J, Stokes MG, Harrison PJ, et al. Resting GABA and glutamate concentrations do not predict visual gamma frequency or amplitude. Proc Natl Acad Sci USA. 2014 Jun 24;111(25):9301–6.

128. Hall SD, Barnes GR, Furlong PL, Seri S, Hillebrand A. Neuronal network pharmacodynamics of GABAergic modulation in the human cortex determined using pharmaco-magnetoencephalography. Hum Brain Mapp. 2010 Apr;31(4):581–94.

129. Routley BC, Singh KD, Hamandi K, Muthukumaraswamy SD. The effects of AMPA receptor blockade on resting magnetoencephalography recordings. J Psychopharmacol (Oxford). 2017 Dec;31(12):1527–36.

130. Wyss C, Tse DHY, Kometer M, Dammers J, Achermann R, Shah NJ, et al. GABA metabolism and its role in gamma-band oscillatory activity during auditory processing: An MRS and EEG study. Hum Brain Mapp. 2017 Aug;38(8):3975–87.

131. Arazi A, Toso A, Grent-’t-Jong T, Uhlhaas PJ, Donner TH. Large-scale maps of altered cortical dynamics in early-stage psychosis are related to GABAergic and glutamatergic neurotransmission. Sci Adv. 2025 Aug 15;11(33):eads0400.

132. Ros T, Kwiek J, Andriot T, Michela A, Vuilleumier P, Garibotto V, et al. PET imaging of dopamine neurotransmission during EEG neurofeedback. Front Physiol. 2020;11:590503.

133. Melgari J-M, Curcio G, Mastrolilli F, Salomone G, Trotta L, Tombini M, et al. Alpha and beta EEG power reflects L-dopa acute administration in parkinsonian patients. Front Aging Neurosci. 2014 Nov 5;6:302.

134. Mackintosh AJ, de Bock R, Lim Z, Trulley V-N, Schmidt A, Borgwardt S, et al. Psychotic disorders, dopaminergic agents and EEG/MEG resting-state functional connectivity: A systematic review. Neurosci Biobehav Rev. 2021 Jan;120:354–71.

135. Bosboom JLW, Stoffers D, Stam CJ, Berendse HW, Wolters EC. Cholinergic modulation of MEG resting-state oscillatory activity in Parkinson’s disease related dementia. Clin Neurophysiol. 2009 May;120(5):910–5.

136. Wiesman AI, Vinding MC, Tsitsi P, Svenningsson P, Waldthaler J, Lundqvist D. Cortical effects of dopamine replacement account for clinical response variability in parkinson’s disease. Mov Disord. 2025 Aug;40(8):1551–60.

137. Deco G, Sanz Perl Y, Vohryzek J, Luppi AI, Kringelbach ML. Neurotransmission-modulated whole-brain computation captures full task repertoire. Cell Rep. 2026 Jan 27;45(1):116816.

138. Oostenveld R, Fries P, Maris E, Schoffelen J-M. FieldTrip: Open source software for advanced analysis of MEG, EEG, and invasive electrophysiological data. Comput Intell Neurosci. 2011;2011:156869.

139. Marquetand J, Vannoni S, Carboni M, Li Hegner Y, Stier C, Braun C, et al. Reliability of Magnetoencephalography and High-Density Electroencephalography Resting-State Functional Connectivity Metrics. Brain Connect. 2019 Sep;9(7):539–53.

140. Garcés P, López-Sanz D, Maestú F, Pereda E. Choice of Magnetometers and Gradiometers after Signal Space Separation. Sensors. 2017 Dec 16;17(12).

141. Saad ZS, Reynolds RC. SUMA. Neuroimage. 2012 Aug 15;62(2):768–73.

142. Box GEP, Jenkins GM, Reinsel GC, editors. Time Series Analysis: Forecasting and Control. 3rd ed. Upper Saddle River, NJ: Prentice Hall; 1994.

143. Merker B. Silver staining of cell bodies by means of physical development. J Neurosci Methods. 1983 Nov;9(3):235–41.

144. Van Essen DC, Ugurbil K, Auerbach E, Barch D, Behrens TEJ, Bucholz R, et al. The Human Connectome Project: a data acquisition perspective. Neuroimage. 2012 Oct 1;62(4):2222–31.

145. Scheinost D, Noble S, Horien C, Greene AS, Lake EM, Salehi M, et al. Ten simple rules for predictive modeling of individual differences in neuroimaging. Neuroimage. 2019 Jun;193:35–45.

